# Exosomes Released from Senescent Cells and Circulatory Exosomes Isolated from Human Plasma Reveal Aging-associated Proteomic and Lipid Signatures

**DOI:** 10.1101/2024.06.22.600215

**Authors:** Sandip Kumar Patel, Joanna Bons, Jacob P. Rose, Jessie R. Chappel, Rebecca L. Beres, Mark A. Watson, Corey Webster, Jordan B. Burton, Roland Bruderer, Pierre-Yves Desprez, Lukas Reiter, Judith Campisi, Erin S. Baker, Birgit Schilling

**Affiliations:** The Buck Institute for Research on Aging, Novato, CA 94945, USA; Bioinformatics Research Center, Department of Biological Sciences, North Carolina State University, Raleigh, NC 27606, USA; Department of Chemistry, University of North Carolina at Chapel Hill, NC 27514, USA; Biognosys AG, 8952 Schlieren, Switzerland; California Pacific Medical Centre, Research Institute, San Francisco, CA 94107, USA

**Keywords:** Proteomics, senescence, exosome senescence-associated secretory phenotype (SASP), data-independent acquisitions, aging

## Abstract

Senescence emerged as significant mechanism of aging and age-related diseases, offering an attractive target for clinical interventions. Senescent cells release a senescence-associated secretory phenotype (SASP), including exosomes that may act as signal transducers between distal tissues, propagating secondary or bystander senescence and signaling throughout the body. However, the composition of exosome SASP remains underexplored, presenting an opportunity for novel unbiased discovery. We present a detailed proteomic and lipidomic analysis of exosome SASP using mass spectrometry from human plasma from young and older individuals and from tissue culture of senescent primary human lung fibroblasts. We identified ∼1,300 exosome proteins released by senescent cells induced by three different senescence inducers. In parallel, a human plasma cohort from young (20–26 years) and old (65–74 years) individuals revealed over 1,350 exosome proteins and 171 plasma exosome proteins were regulated when comparing old vs young individuals. Of the age-regulated plasma exosome proteins, we observed 52 exosome SASP factors that were also regulated in exosomes from the senescent fibroblasts, including serine protease inhibitors (SERPINs), Prothrombin, Coagulation factor V, Plasminogen, and Reelin. 247 lipids were identified in exosome samples. Following senescence induction, identified phosphatidylcholines, phosphatidylethanolamines, and sphingomyelins increased significantly indicating cellular membrane changes. Interestingly, significantly changed proteins were related to extracellular matrix remodeling and inflammation, both potentially detrimental pathways that can damage surrounding tissues and even induce secondary senescence. Our findings reveal mechanistic insights and potential senescence biomarkers, enabling a better approach to surveilling the senescence burden in the aging population and offering therapeutic targets for interventions.

## Introduction

The World Health Organization (WHO) estimates that one out of every six individuals in the world will be aged 60 or older by 2030 [1]. This demographic shift will have far-reaching social, economic, and healthcare implications. The aging population is at an increased risk for chronic diseases, which could result in a significant public health crisis [2, 3]. Therefore, there is a critical need to prioritize healthy aging, which begins with comprehending the fundamental molecular basis of aging to identify potential interventions and biomarkers.

Cellular senescence is considered a crucial mechanism of aging because senescent cells accumulate in different tissues over time and contribute to the aging process [4]. Senescence is the process by which cells become cell cycle arrested while still staying viable, metabolically active, and undergoing distinct phenotypic alterations. Senescence can be induced by various intrinsic factors, such as telomer shortening (Hayflick’s limit) [5], by extrinsic factors, such as irradiation (IR) and chemical reagents (e.g., doxorubicin, doxo), or via Mitochondrial Dysfunction-Associated Senescence (MiDAS) [6, 7]. The various senescence inducers all exhibit varied modes of action, for example, irradiation causes double-stranded breaks in DNA [8], doxorubicin induces DNA cross-linking [9], while disruption in normal mitochondrial function results in MiDAS [6]. Senescent cells exert their pleiotropic biological effects by secretion of senescence-associated secretory phenotype (SASP) [10–12]. The SASP represents a myriad of pro-inflammatory molecules, growth factors, extracellular matrix (ECM) metalloproteinases, microRNAs, and lipids that are either directly secreted or packaged into exosomes [13–15].

Exosomes are small, lipid-bilayer enclosed nanoparticles derived from cells with a diameter ranging between 30 nm and 150 nm. They are known to transport various biomolecules to recipient cells in a paracrine or endocrine manner, allowing them to act as signal transducers between distal tissues [16]. The cargo carried by circulating exosomes includes proteins, lipids, miRNA and other components, which all have been implicated in aging and age-related diseases [17]. However, the comprehensive characterization of exosome SASP and their role in potentially propagating senescence (secondary/bystander senescence) and how they may influence the aging progression is still largely understudied. Several challenges include sub-optimal enrichment strategies that are compatible with mass spectrometry technologies and protein contamination from biofluids, which often impede downstream omics processing [18].

Current methods for isolating exosomes commonly rely on physical properties of extracellular vesicles (EVs) or target-specific surface markers [19]. A novel approach, Mag-Net, enriches extracellular vesicles from plasma through electrostatic interactions, providing a promising alternative [20]. Ultracentrifugation is one of the more common methods for exosome enrichment [21]. However, this low-throughput method can lead to contamination from other EVs, high molecular weight proteins, and highly abundant proteins. To enhance exosome purity, size exclusion chromatography (SEC) is often used before centrifugation or filtration [22]. Developing high-throughput workflows for identifying and quantifying circulating exosome SASP components (secreted from various senescent cell types) in plasma is a valuable resource for characterizing cellular senescence signatures by identifying (i) specific affected tissues, (ii) biomarkers to measure biological age non-invasively, and (iii) novel targets for anti-aging therapeutics.

In this study, we developed a platform to enrich exosomes from plasma and tissue culture for the analysis of exosomal proteins, lipids, and miRNA (Fig. 1). Additionally, we generated a novel “human exosome protein spectral library” using data-dependent acquisition (DDA) and performed data-independent acquisition (DIA) workflows for exosome protein quantification. This development introduces quantitative plasma exosome workflows, where DIA-MS acquisitions [23, 24] are searched efficiently in Spectronaut (Biognosys). Our study showed that senescent fibroblasts induced by three stimuli (IR, doxo, and Antimycin A-induced MiDAS) released highly complex and dynamic overlapping exosome profiles, some of which overlap with profiles we also observed in the human plasma exosomes from older individuals. Importantly, in exosomes from senescent cells we identified many categories of significantly changing proteins related to extracellular matrix remodeling and inflammation, which are both implicated in accelerating secondary senescence. Our study aims to address several critical knowledge gaps in the field of exosome-associated SASP, including identifying aging-specific exosome signatures, cell-specific contributions, and local versus systemic effects. We also explore molecular pathways through which exosomes propagate senescence, inflammation, and tissue remodeling. To advance this understanding, we optimized exosome isolation workflows from both cell culture and plasma, enabling robust multi-omics analyses. Our study has identified and quantified exosome cargo, including not only proteins but also significantly altered lipids and miRNAs. These findings enhance our understanding of exosome composition in aging and provide a foundation for developing senescence-targeted diagnostics and therapeutics to manage age-associated diseases and promote healthy aging.

**Fig. 1:**
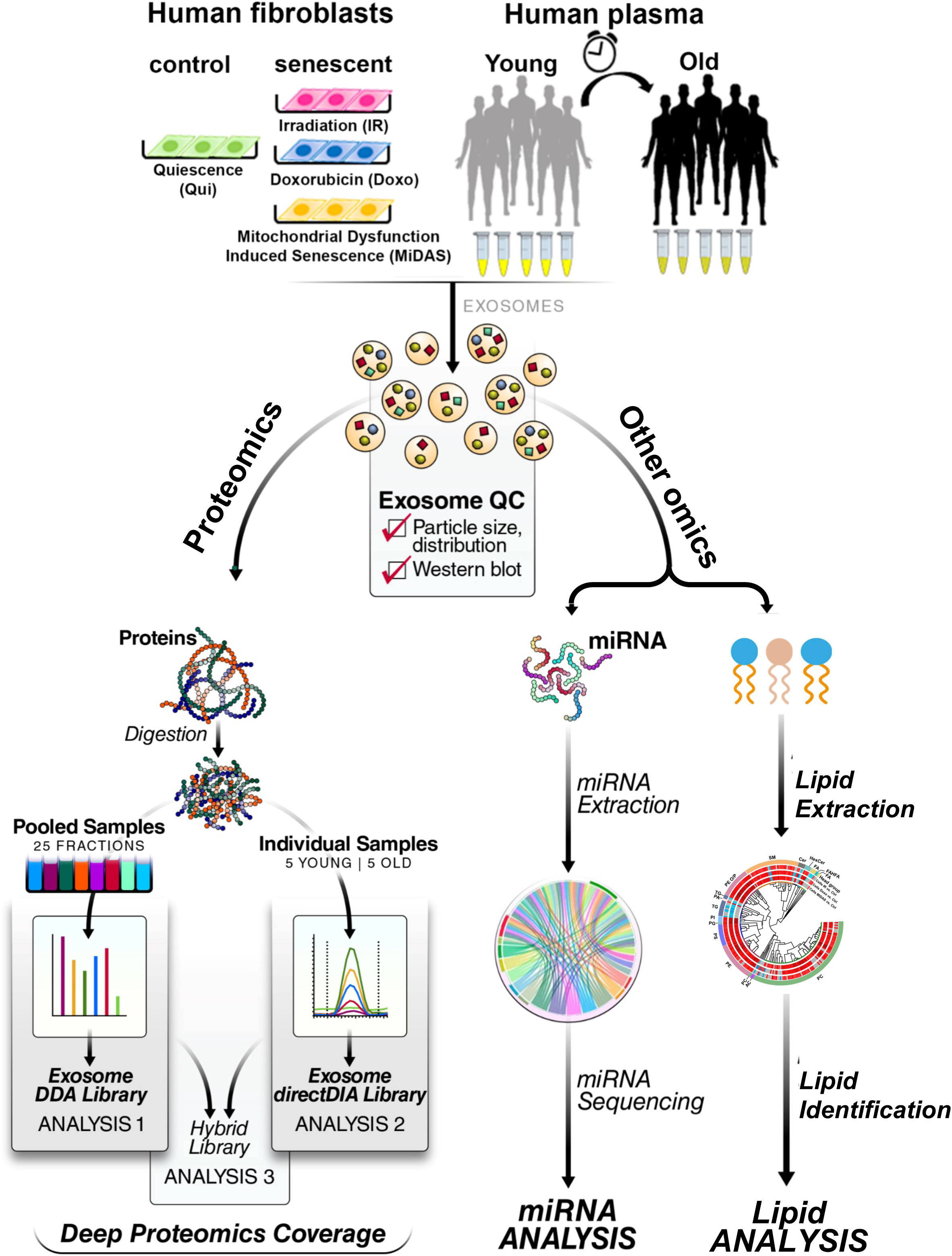
Multi-omics workflow for aging/senescence exosome biomarker discovery. **A.** Two different sample types, primary human lung fibroblasts and human plasma, were investigated. Three senescence stimuli, Irradiation; IR, Doxorubicin; doxo, and Antimycin A-induced mitochondrial dysfunction-associated senescence; MiDAS, were used to generate senescent primary human lung fibroblasts (n=4, each condition). Quiescent cells (Qui) or DMSO-treated fibroblasts were used as control. Plasma from young (20-26 years old) and older (65-74 years old) individuals was used (n=5 each). Exosomes were enriched by sequential size-exclusion chromatography and ultrafiltration (SEC/UF). Immunoblotting and particle size analysis was conducted to confirm enriched, high-quality, and intact exosomes. Three different protein spectral libraries; DDA, directDIA, and hybrid, were generated. Multicomponent Senescence/Aging Biomarkers were identified by high-throughput quantitative exosome multi-omics (proteomics, lipidomics, and miRNA) analysis.

## Results

### Exosome enrichment workflow from a plasma aging cohort and from primary cultured senescent cells

We performed a comprehensive proteomic analysis of exosome SASP from senescent primary human lung fibroblast cells (IMR90) induced by three different stimuli, irradiation (IR), doxorubicin (doxo), and Antimycin A-induced MiDAS. Furthermore, we profiled the proteome and miRNA contained in exosomes from plasma from young (20-26 years old) and older (65-74 years old) individuals (Fig.1).

To improve the purity of exosome enrichment for downstream processing we developed a robust workflow for exosome enrichment using sequential size-exclusion chromatography (SEC) followed by ultrafiltration (SEC/UF) or SEC followed by ultracentrifugation (SEC/UC; Fig. S1a). Both SEC/UC and SEC/UF yielded equivalent protein groups identification for exosomes by mass spectrometric proteomics analysis (Fig. S1b). With consideration for ease, time, and cost-effectiveness, we proceeded with SEC/UF for downstream proteomics and miRNA analysis (Fig. 2A). The details of SEC elution and pooling strategies are provided in Fig. 2B. Twenty SEC-extracted plasma fractions (1 mL each) were initially analyzed by spectrophotometry as a crude assessment of exosome enrichment (15-35 mL in the fractionation scheme). The majority of plasma exosomes were eluted within the first 10 fractions (*i.e.* 10 mL), whereas contaminating plasma proteins eluted more slowly, with a maximum protein concentration achieved by the 20^th^ fraction (Fig. 2C). A pool of all the fractions containing exosomes were estimated to have ∼1×10^10^ exosome particles per mL of plasma with a qNano Gold nanoparticle characterization instrument, which uses tunable resistive pulse sensing to measure particles (Fig. 2D). For immunoblotting, from the 10 mL containing exosomes (Fig. 2C) we pooled each two consecutive fractions into one, resulting in a total of five exosome-enriched fractions (E1–E5), one post-EV fraction (P1), and a pooled soluble protein fraction (P10) containing all subsequent fractions (Fig. 2B). Fractions E2–E5 stained positively for the exosome-specific protein markers CD9 (27 kDa) and Tumour Susceptibility Gene 101 (TSG101; 44 kDa; Fig. 2E-F). Using the SEC/UF workflow, we observed more intense bands for exosome-specific markers and less contaminating bands from protein contaminants IgG and albumin (which pose challenges in downstream proteomic analysis) compared to UC (Fig. 2E-F). These results highlighted our rational to proceed with the SEC/UF approach. Correspondingly, we also performed exosome enrichment and subsequent exosome quality control (QC) for exosomes from human primary fibroblasts IMR90 (Fig. S1c). Overall, our SEC/UF workflow led to highly enriched, intact, and ‘contaminant-free’ (certainly contaminant-reduced) exosomes from both isolation schemes either from human plasma as well as from human primary fibroblasts. The biological outcomes and pathways are discussed in the below sections.

**Fig. 2:**
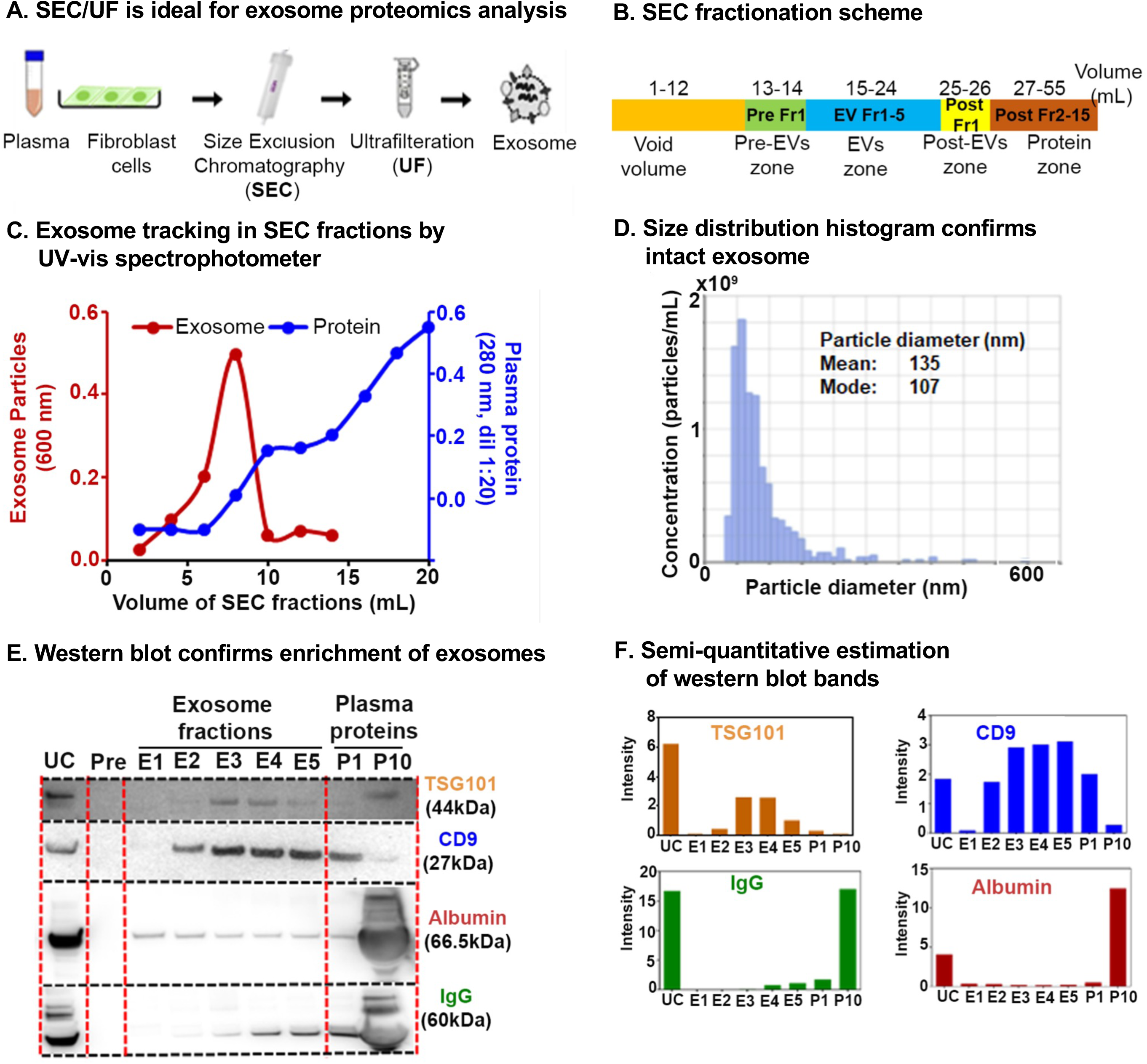
Exosome enrichment from plasma and primary human lung fibroblasts (IMR90) and characterization. **A.** Exosome isolation workflow using sequential size-exclusion chromatography and ultrafiltration (SEC/UF). **B.** Scheme for SEC fraction collection. **C.** Spectrophotometric quantification of exosomes in SEC fractions. Red, Exosome (OD_600_); Blue, plasma proteins (OD_280_, dilution 1:20). **D.** Exosome size distribution analysis by qNano Gold. Mean diameter; 135 nm, Mode diameter; 107 nm. **E.** Immunoblot of exosome protein markers; CD9 antigen (CD9) and Tumor susceptibility gene 101 protein (TSG101) in different exosome fractions to confirm enrichment. IgG and albumin were used as determinants of plasma protein contaminants. E1–E5: exosome fractions, P1: the pool of fractions 11 + 12, and P10: the pool of fractions 29 + 30 (1:20 dilution) were loaded for comparison between plasma proteins and exosome fractions. **F.** Semi-quantitative estimation of the volume intensities of western blot bands using Image J software.

### Mass spectrometric quantification of exosome protein cargo via label-free data-independent acquisition (DIA)

Data-independent acquisition (DIA)-mass spectrometry (MS) provides comprehensive and highly quantitative workflows for protein identification analyzing complex biological matrices, such as plasma, but also from more specialized ‘sub-compartments’, such as exosomes and/or extracellular vehicles. DIA-MS [23–25] approaches have tremendously enhanced protein coverage, deep protein profiling and quantitative precision by employing an unbiased MS/MS data acquisition of precursor ions, leading to confident and reproducible peptide identification and quantification across biological replicates. To optimally search using mass spectrometric spectral libraries for data processing, we developed custom, in-house spectral libraries for human plasma-exosome samples from five young (20–26 years) and five older (65-74 years) individuals (Table S1), using two different data acquisition approaches (Fig. 1). The first approach using data-dependent acquisition (DDA) built a comprehensive exosome spectral library from highly fractionated samples, that served as a resource for future work, and that can be applied to any future DIA-MS analyses of human exosome samples. More specifically, we generated a DDA spectral library from DDA acquisitions of 25 generated reversed-phase fractions from pooled plasma exosome proteins. A total of 2,323 protein groups was identified upon processing with the Spectronaut (Biognosys) search algorithm (Fig. S2A; Table S2A), among which 1,902 protein groups were identified with at least two unique peptides (Table S2B). The second approach using data-independent acquisition (DIA) was introduced to feature a robust, precise, and straightforward methodology that does not require an up-front generation of deep spectral libraries by fractionation, which can be time-consuming, labor-intensive, and typically requires a larger amount of starting material. Plasma exosome samples from 10 human individuals were subjected to DIA-MS analysis and subsequently processed using the directDIA algorithm within Spectronaut. We also combined the DDA and directDIA spectral libraries to develop a hybrid library consisting of overall 2,225 protein groups (Fig. S2A; Table S2E), of which 1,808 proteins were identified with at least two unique peptides (Table S2F). The hybrid spectral library was used to search the plasma exosome DIA-MS data reported in this study (Table S2F). For more details regarding the spectral library approaches and additional metrics and quality control, as well as comparisons with other publicly available biofluid exosome datasets (urine exosomes [26]), please see Figure S2 and Table S2A-H. Our generated hybrid spectral library showed an overlap of 79% with the Plasma Exocarta database [27], confirming a highly enriched and robust isolated exosome proteins. Interestingly, 370 (∼21%) plasma exosome proteins were uniquely identified in our study (Fig. S2H), potentially representing novel plasma exosome proteins.

### Distinct exosome SASP protein profiles obtained from senescent human primary lung fibroblasts exposed to different senescence inducers

To evaluate the impact of different senescence stimuli on exosome SASP proteins, we induced senescence in human primary fibroblasts by (a) Irradiation (IR), (b) doxorubicin (doxo), and (c) Antimycin A (MiDAS) (Fig. 3A (i)). Either quiescent (Qui) or DMSO-treated (DMSO) fibroblasts were used as a control. Each senescence stimulus and the control group comprised four biological replicates. Fig. 3A (ii) provides a detailed account of the exosome SASP collection following senescence induction. Briefly, we induced senescence in fibroblasts by IR, doxo, or MiDAS and allowed 10 days for the senescent phenotype to develop as previously described [10]. In parallel, the control cells were brought into quiescence state (Qui) by incubating in 0.2% serum (minimal media) for 3 days by mock irradiation or treatment with DMSO in 0.2% serum for 1 day. Senescence-induced cells and control cells (Qui or DMSO) were cultured in serum-free medium for 1 day before collection of the conditioned media containing exosome SASP for the mass spectrometric end-point analysis. Cell viability assays confirmed no detectable difference in apoptosis/cell death in senescent and control fibroblast cells measured by SYTOX™ Green Nucleic Acid Stain (Fig. S3A). Furthermore, EdU and SA-β-Gal counter staining confirmed senescence (more than 90%) in IR-, doxo-, and MiDAS-induced senescent IMR90 cells compared to Qui control, while ∼10% of senescent and Qui controls were dividing cells (Fig. 3B). In MiDAS, mitochondrial dysfunction was observed by a decrease in NAD+/NADH ratio indicating the electron transport chain was inhibited (Fig. S3B).

**Fig. 3:**
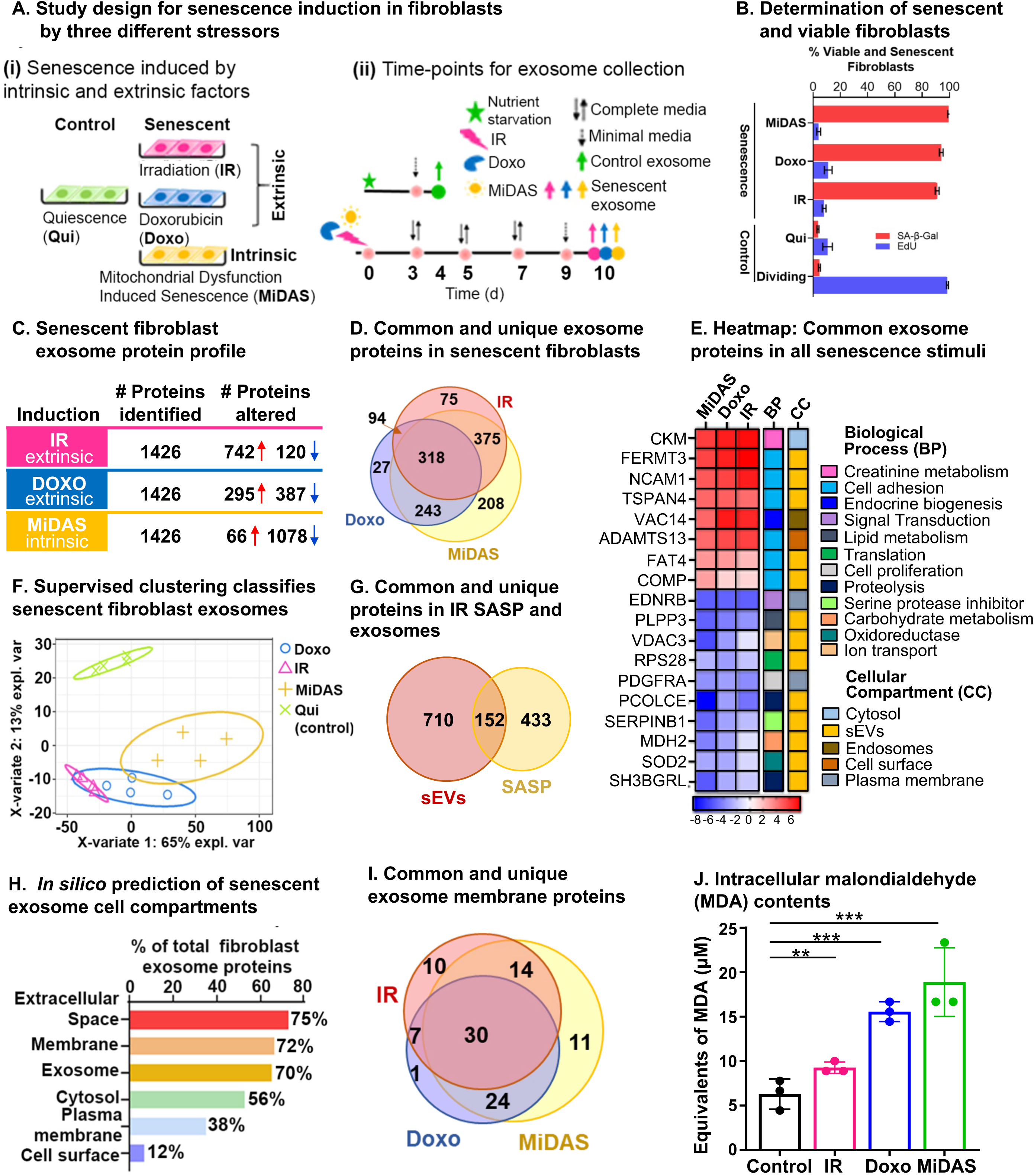
Senescence stimuli specific exosome protein signatures in primary human lung fibroblasts (IMR90). **A (i).** Study design showing three different senescence stimuli, Irradiation; IR, Doxorubicin; doxo, and Antimycin A-induced mitochondrial dysfunction-associated senescence; MiDAS, were used to generate senescent primary human lung fibroblasts (n=4, each condition), and Quiescent cells (Qui) were used as control. **(ii)** A timeline indicating treatment days and endpoints for the three different senescence inducers and control conditions. **B.** Bar graph showing the percentage of viable and senescent fibroblasts in the different senescence-induction conditions using Edu (shown in blue) and SA-β-Gal (shown in red). **C.** Summary of significantly altered exosome proteins (q-value < 0.05 and ≥ 1.5-fold change) in senescent human lung fibroblasts compared to quiescent cells following IR, doxo, and MiDAS senescence induction. **D.** Venn diagram showing overlapping and unique protein signatures of senescent human lung fibroblast exosomes induced by IR, doxo, and MiDAS. **E.** Heatmap displaying the exosome protein fold change averaged across replicates, showing the top 18 altered proteins, increased (red) and decreased (blue) due to senescence. The intensities averaged for Qui were used as the baseline. ClueGO biological pathways and cell compartments are presented. BP; Biological process, CC; Cellular process. **F.** Partial least squares-discriminant analysis (PLS-DA) clustered senescent fibroblasts from Quiescent control. The two variates explaining the most significant variations are shown. **G.** Venn diagram showing overlapping and unique exosome SASP (sEVs) and soluble SASP protein signatures of IR-induced senescent fibroblasts. **H.** Enrichment analysis of Gene Ontology/cellular compartments overrepresented among protein contents of senescent fibroblast exosomes. **I.** Venn diagram showing overlapping and unique surface protein signatures of senescent fibroblast exosomes induced by IR, doxo, and MiDAS. **J.** Bar graph showing malondialdehyde cellular content, a product of lipid peroxidation used as a marker for intracellular oxidative stress measured in the different senescence stimuli.

Size exclusion/ultrafiltration (SEC/UF)-enriched exosome proteins were extracted and processed for label-free data-independent acquisition (DIA). The DIA-MS approach facilitated sensitive and accurate quantification of exosome SASP proteins by quantification of the tandem mass spectrometry (MS2) fragment ions, specifically quantifying the ‘peak area under the curve’ from extracted ion chromatograms (XICs). We quantitatively compared exosome proteins (≥ 2 unique peptides) released by senescent fibroblasts with exosome proteins from Qui controls, and we cataloged significantly changed proteins (q-value ≤ 0.05) that featured an absolute fold change ≥ 1.5-fold (comparing senescent vs control) (Fig. 3C; Table S3A-F). Exosome proteins were released at considerably higher levels by senescent fibroblasts compared to quiescent control cells, and we refer to them as ‘exosome SASP’. Our unbiased proteomic profiling identified 1,426 exosome proteins amongst the three different senescent inducers, a large fraction of which 60%, 48%, and 80% were significantly up- or down-regulated in IR, doxo, or MiDAS, respectively (Fig. 3C). Interestingly, most of the significantly changed exosome proteins (742) were markedly upregulated in the IR condition; in contrast, most exosome proteins (1,078) were down-regulated in MiDAS-induced senescence (Fig. 3C). To visualize the significant changes, the volcano plots of senescent fibroblasts versus Qui control (senescence/Qui) showed the significantly altered protein levels (q-value ≤ 0.05, |log_2_fold change| ≥ 0.58) (Fig. S3c, Table S3b, d, f). Up-regulated proteins in ‘IR vs Qui’ included Fermitin family homolog 3 (FERMT3), Creatine kinase M-type (CKM), Myosin-9 (MYH9) as well as numerous ribosomal proteins (RPLs and RPs) and eukaryotic translation initiation factors (EIFs), while ECM proteins collagens, laminins, Decorin (DCN), Biglycan (BGN), and nidogens were down-regulated. With respect to ‘Doxo vs Qui’, up-regulation of several proteasome subunits (PSMCs, PSMDs, PSME2), Protein VAC14 homolog (VAC14), and Collagen alpha-1(XII) chain (COL12A1), and down-regulation of Endothelin receptor type B (EDNRB), Phospholipid phosphatase 3 (PLPP3), Neuroblast differentiation-associated protein AHNAK were observed. ‘MiDAS vs Qui’ comparison featured up-regulated Neutral amino acid transporter A (SLC1A4), complement proteins and fibrinogens, and down-regulated Pantetheinase (VNN1), Procollagen C-endopeptidase enhancer 1 (PCOLCE), collagens, and laminins. Although we discovered distinct protein signatures per each of the senescence induction with proteins that are uniquely up- and down-regulated with senescence, we also observed 318 exosome proteins that are common between the different inductions (Fig. 3D). Specifically, we identified 46 up- and 103 down-regulated exosome proteins that are common among the different senescence stimuli (Table S3G and S3H, respectively). Top common up-regulated SASP proteins were Fermitin family homolog 3 (FERM3), Creatine kinase M-type (CKM), Protein VAC14 homolog (VAC14), Neural cell adhesion molecule 1 (NCAM1), and Tetraspanin-4 (TPS4). Top common down-regulated proteins upon senescence induction included Procollagen C-endopeptidase enhancer 1 (PCOLCE), Endothelin receptor type B (EDNRB), Phospholipid phosphatase 3 (PLPP3), Leukocyte elastase inhibitor (SERPINB1), and mitochondrial Superoxide dismutase [Mn] (SOD2). The top up- and down-regulated exosome SASP proteins common between MiDAS, doxo, and IR conditions are grouped into 12 biological processes (BP), such as cell adhesion, proteolysis, and metabolism, and 5 cellular compartments (CC) (Fig. 3E); 72% of these common proteins are known exosome (or EV) proteins. The clustering by partial least squares-discriminant analysis (PLS-DA) confirmed distinct exosome profile clustering between senescent fibroblasts and quiescent control fibroblasts, whereby clustering of senescent conditions showed overlap in their exosome profiles (Fig. 3F). Furthermore, we compared our IR-induced senescent fibroblast exosome proteins with the IR-induced soluble SASP from our previous study by Basisty and Schilling et al. [10] and found 152 overlapping proteins, while 710 proteins were unique to the exosomes (EVs; Fig. 3G, Table S3I). Among the 152 common proteins were identified Myosin-9 (MYH9), Growth/differentiation factor 15 (GDF15), Interstitial collagenase (MMP1), C-type lectin domain family 11 member A (CLEC11A), and Metalloproteinase inhibitor 1 (TIMP1), which were reported to associate with several traits of aging and constitute a proteomic signature of aging [28, 29]. Exosome-specific proteins included Clusterin (CLU), Serpin B5, dynactin subunits (DCTNs), numerous proteins related to DNA and RNA metabolism (RNA helicase DDXs, heterogeneous nuclear ribonucleoproteins HNRNPs, EIFs, RPs), proteasome subunits, and vacuolar protein sorting-associated proteins (VPSs). The significantly altered senescent fibroblast exosome proteins identified in the study were categorized based on subcellular locations: extracellular space (75%), membrane (72%), exosome (70%), cytosol (56%), plasma membrane (38%), and cell surface (12%) (Fig. 3H). *In silico* analysis of the significantly altered senescent exosome proteins identified 97 out of 1,340 proteins as surface proteins, of which 30 were consistent in all senescence stimuli (Fig. 3I, Table S3J) and included Neural cell adhesion molecule 1 (NCAM1), A disintegrin and metalloproteinase with thrombospondin motifs 13 (ADAMTS13), Neutral amino acid transporter A (SLC1A4), and Tumor necrosis factor ligand superfamily member 4 (TNFSF4). These upregulated senescent surface proteins have the potential to be developed into aging biomarkers. Our proteomics data indicated a decrease in antioxidant proteins in senescent exosomes compared to Qui controls, which is consistent with the higher intracellular oxidative stress in senescent fibroblasts (Fig. 3J). Interestingly, MiDAS induction showed the most elevated cellular oxidative stress, followed by doxorubicin (doxo) and irradiation (IR) as shown in Fig. 3J.

### Plasma exosomes isolated from an aged human cohort display unique protein markers

We performed a pilot study using human plasma samples from five young (20–26 years) and five older individuals (65-74 years) (Table S1) to determine exosome-specific age signatures. We used DIA-MS protein identification and quantification to analyze plasma exosomes from young and elderly individuals and processed the data using our custom hybrid spectral library (generated as described above) (Fig. S4). We reproducibly identified and quantified a total of 1,356 protein groups (≥ 2 unique peptides) (Table S4B)

Distinct and separate clustering of protein profiles from plasma exosomes from young and old individuals were revealed by PLS-DA (Fig. 4A). Volcano plots of old versus young (old/young) individuals showed 171 significantly altered exosome proteins (q-value ≤ 0.05, |log_2_fold change| ≥ 0.58), of which 117 were down-regulated, whereas 54 were up-regulated in the older individuals (Fig. 4B; Table S4C). A selection of key importance up- and down-regulated aging plasma exosome proteins (26) are shown in Fig. 4C. Top up-regulated proteins included LIM and SH3 domain protein 1 (LASP1), Hemopexin (HPX), Eosinophil peroxidase (EPX), Band 3 anion transport protein (SLC4A1), Leucine-rich alpha-2-glycoprotein (LRG1), as well as several Serpins. Top down-regulated proteins were Calmodulin-like protein 5 (CALML5), Deoxyribose-phosphate aldolase (DERA), Peroxidasin homolog (PXDN), Aminopeptidase N (ANPEP), and Apolipoprotein L1 (APOL1). Out of them, 80% are classified as exosome proteins, demonstrating the high yield and purity of the extracted plasma exosomes, and span nine different biological processes, such as exocytosis, immune response, signal transduction, and proteolysis (Fig. 4C).

**Fig. 4:**
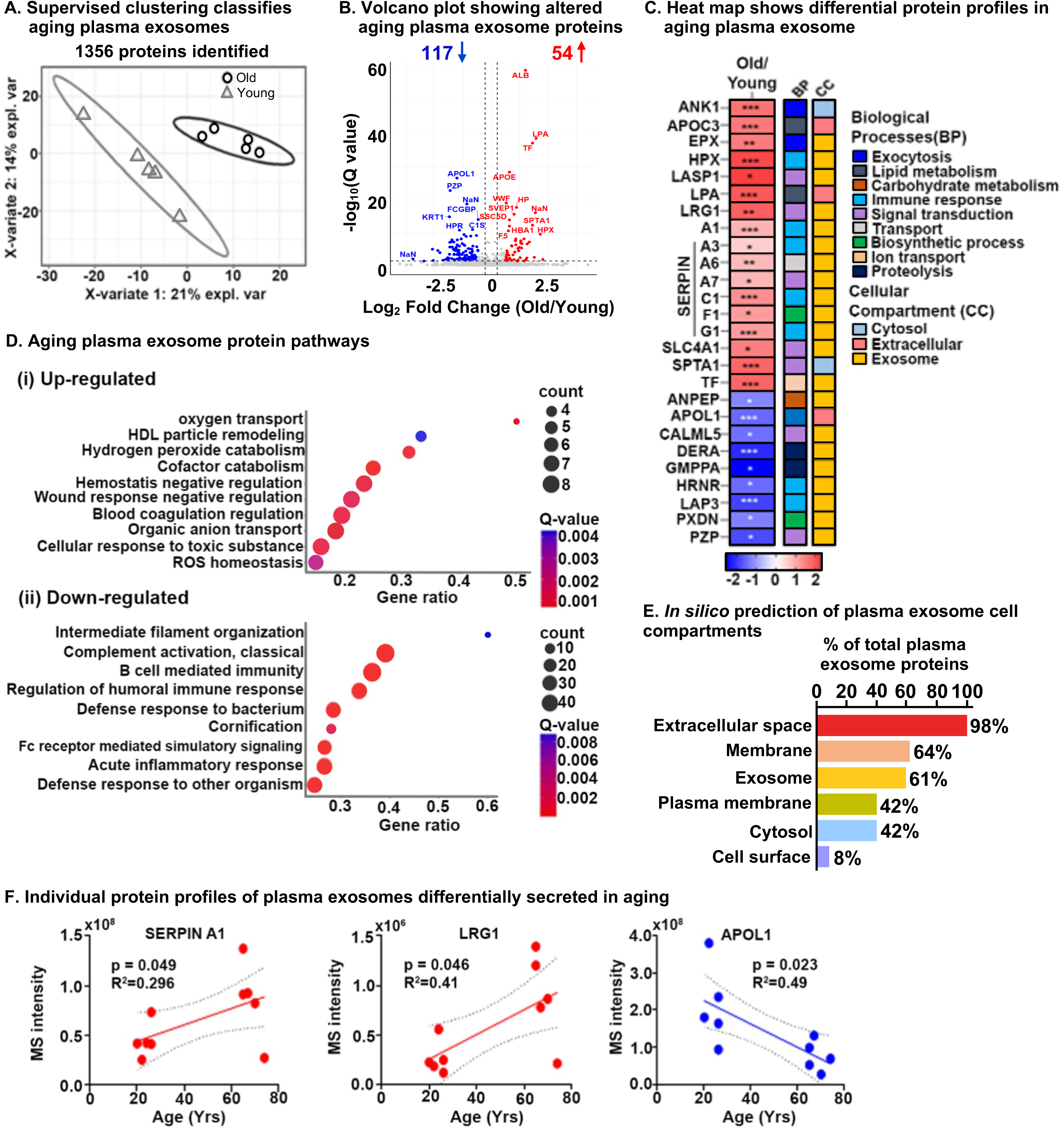
Age-specific plasma exosome protein signatures in old and young cohorts. **A.** Partial least squares-discriminant analysis (PLS-DA) clustered old and young cohorts based on 1356 protein groups. The two variates explaining the most significant variations are shown. **B.** Volcano plot showing significantly altered age-specific plasma exosome proteins (q-value < 0.05 and ≥ 1.5-fold). Red, upregulated; Blue, downregulated; and Gray, No significant change. A few selected differentially abundant proteins are labeled. **C.** Heatmap displaying the plasma exosome protein fold change averaged across replicates, showing the top 26 altered proteins increased (red) and decreased (blue) due to Aging. The intensities averaged for young plasma exosomes were used as the baseline. ClueGO biological pathways and cell compartments are presented. BP; Biological process, CC; Cellular process. P-value * ≤ 0.05, ** ≤ 0.01, *** ≤ 0.001. **D.** The most significant age-related exosome protein pathways are (i) upregulated and (ii) downregulated. **E.** Enrichment analysis of gene-ontology/cellular compartments overrepresented among aging exosome proteins. **F.** Three examples of individual protein MS intensities and how the intensity changes with age.

To identify the biological relevance of our results, the differentially changing exosome proteins in aging plasma from our DIA-MS analysis were integrated and highlighted using ConsensusPathDB-human metabolic pathways and the Gene Ontology (GO) annotation toolbox. Pathway enrichment analysis indicated remodeling in the plasma lipoprotein particles, changes in blood coagulation, negative regulation of response to wounding, and reactive oxidant activity – these pathways were significantly upregulated in plasma exosomes from older individuals. Conversely, defense response to bacteria, acute inflammatory response, humoral immune response regulation, cornification, and intermediate filament cytoskeleton organization were pathways that were significantly down-regulated (Fig. 4D; Table S4J & S4K). The significantly altered plasma exosome proteins identified in the study were categorized based on subcellular locations: extracellular space (98%), membrane (64%), exosome (61%), plasma membrane (42%), cytosol (42%), and cell surface (8%) (Fig. 4E). Individual changes in the exosome protein level of Alpha-1-antitrypsin (SERPINA1) and Leucine-rich alpha-2-glycoprotein (LRG1) positively correlated with older age (upregulation), whereas Apolipoprotein L1 (APOL1) negatively correlated with age (downregulation) as shown in Fig. 4F, which further implies distinct aging signatures in human plasma.

### Significant variations in lipid compositions of exosomes induced by different senescence stimuli

In addition to exosome proteomics, we conducted a parallel lipidomic analysis of exosomes from senescent fibroblasts induced by the three senescence-inducing stimuli. Exosomes enriched by antibodies were subjected to lipidomic analysis using liquid chromatography ion mobility separation collision-induced dissociation mass spectrometry (LC-IMS-CID-MS). By separating lipids based on their LC retention time, IMS collision cross section (which is based on the lipid’s three-dimensional shape), and precursor and fragment ion masses, LC-IMS-CID-MS analyses can reduce ambiguity and increase confidence in lipid identification and quantification. Here, we quantitatively compared lipids from exosomes released from senescent and control fibroblasts and cataloged significantly changed lipids (p_adj_-value ≤ 0.05) that showed a fold change ≥ 2-fold (senescent/control) (Fig. 5A; Table S5A). Lipidomic profiling revealed a substantial number of statistically significant exosome lipids in response to the senescence inductions: with 122 regulated lipids with IR induction, 135 regulated lipids with doxo induction, and 129 regulated lipids with MiDAS induction (yielding a total of 247 changed lipids upon senescence induction considering all inducers). Notably, most of the significantly changed exosome lipids were upregulated in all senescent conditions (Fig. 5A). Although there is variation in the lipids that were up- and downregulated with senescence, 112 were shared amongst all inducers (Table S5B and S5C). Furthermore, distinct clustering of senescent and control fibroblast exosome profiles was revealed by PLS-DA (Fig. 5B). Individual lipid species profiles are further depicted in the heatmap in Fig. 5C. Our results indicate that membrane lipids, such as phosphatidylcholine (PC), phosphatidylethanolamine (PE), phosphatidylserine (PS), sphingomyelin (SM), and hexosylceramide (HexCer) species were significantly upregulated in senescence conditions. Specifically,top up-regulated lipids included PC(16:0_22:4), PC(18:0_24:0), PE(O-18:0/22:5), PE(P-18:0/20:4), PS(18:1_22:0), SM(d18:2/16:0), HexCer(d18:1/24:0). Our analysis also revealed senescence stimuli-specific changes in exosome lipids. Specifically, triglycerides (TG), TG(16:0_18:1_20:4), TG(16:1_18:1_20:4);TG(18:2/18:2/18:2), and TG(54:6), and acylcarnitine (AC) AC(18:0) species were downregulated with the doxo and IR inducers but not with MiDAS-induced senescence (Fig. 5C). Additionally, specific PC and PC plasmalogen species (PC(P-)), PC(P-18:0/20:5) and PC(P-16:0/22:5), were only upregulated with the MiDAS induction (Fig. 5C). Changes in the levels of specific exosome lipids such as PC (18:0_24:0) and SM (d18:2/16:0) are further shown to illustrate they are positively correlated in all three senescent conditions, while TG(54:6) is denoted for its decrease in only the doxo and IR inductions (Fig. 5D), suggesting unique senescence lipid signatures in human fibroblasts. Overall, these significant changes in exosome-specific lipids highlight that they may play a potential role in the senescence process.

**Fig. 5:**
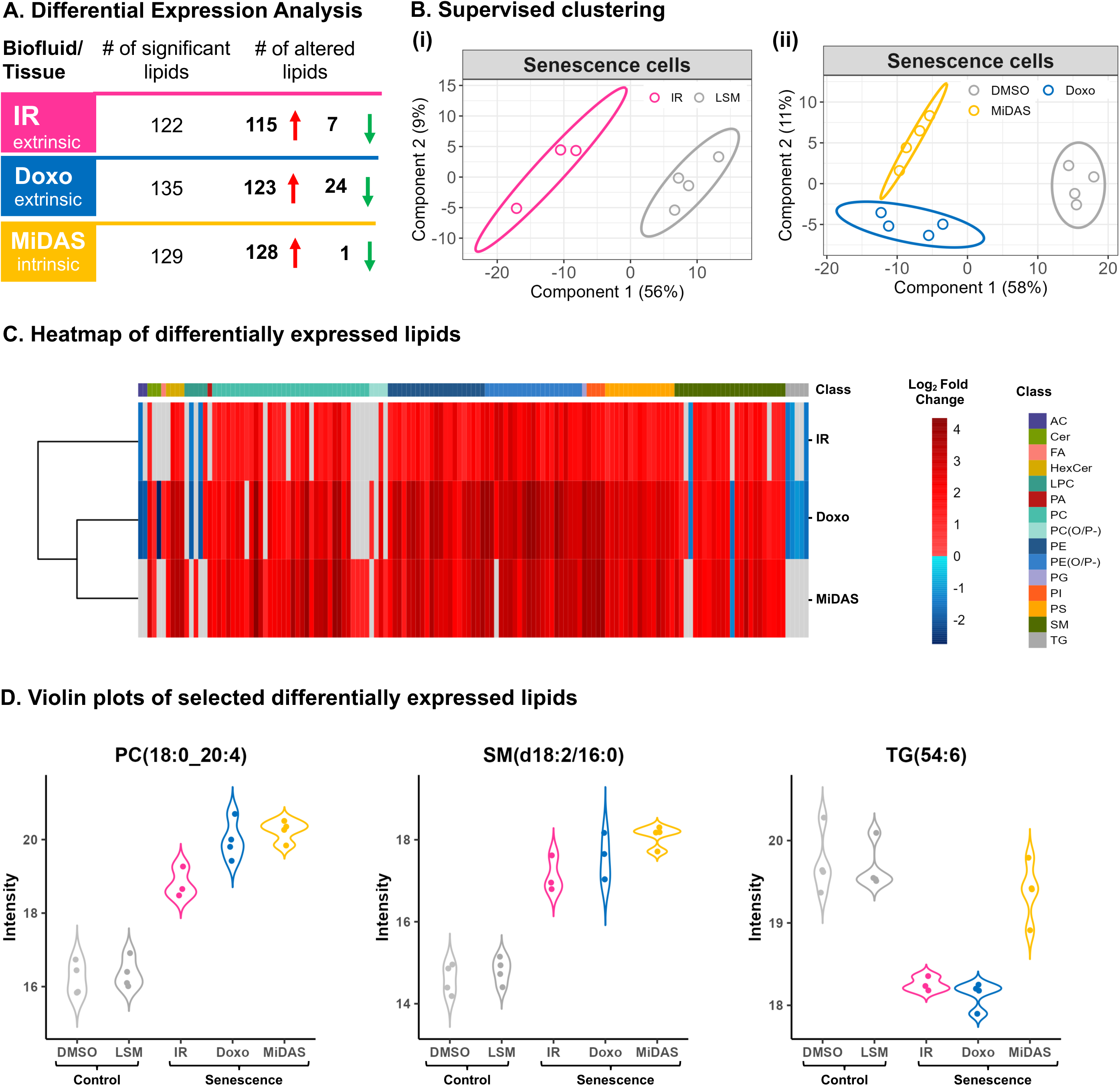
Senescence stimuli specific exosome lipid signatures in primary human lung fibroblasts (IMR90). **A.** Summary of significantly altered exosome lipids (p-value < 0.05 and > 2-fold change) from senescent human lung fibroblasts compared to quiescent cells following IR, doxo, and MiDAS senescence induction. **B.** Partial least squares-discriminant analysis (PLS-DA) clustered senescent human lung fibroblasts from Quiescent (control) based on 247 lipids **(i)** Low serum media (LSM; Qui IR-mock control) vs IR **(ii)** Low serum media (Qui DMSO control) vs doxo and MiDAS. **C.** Heatmap of 145 differentially expressed exosome lipids from 15 different classes. Clustering was performed using Euclidean distance and complete linkage and log_2_ fold change is illustrated to visualize trends with the three senescence inducers. **D.** Violin plots showing three examples of individual lipid profiles where PC(18:0/20:4) and SM(d18:2/16:0) were upregulated with all senescence inducers, while TB(54:6) was downregulated in IR and doxo but not MiDAS, illustrating lipid response differences due to different stimuli. AC: Acylcarnitine, Cer: Ceramide, FA: Fatty acid, HexCer: Hexosylceramide, LPC: Lysophosphatidylcholine, PA: Phosphatidic acid, PC: Phosphatidylcholine, PE: Phosphatidylethanolamine, PG: Phosphatidylglycerol, PI: Phosphatidylinositol, PS: Phosphatidylserine, SM: Sphingomyelin, TG: Triglycerides.

### Exosomes miRNAs are differentially present in human plasma from young and older individuals

To complement the exosome proteomic and lipidomic data, we performed in parallel a pilot analysis of human plasma exosome miRNA cargo to identify differences between young (20–26 years) and older (65-74 years) individuals (Fig. 1). The length of miRNA sequences primarily ranged from 18 to 24 nucleotides (Fig. S5a). In total, 1,375 miRNAs were identified from our SEC/UF-extracted plasma exosomes (Table S6a), of which, 331 miRNA were consistent. 226 out of 331 miRNA were commonly shared between exosomes from both young and older individuals, and 88 and 17 were unique to plasma exosomes from old and young individuals, respectively (Fig. S5b; Table S6b). Volcano plots of old vs. young samples revealed six miRNAs to be significantly altered. hsa-miR-27a-5p, -874-3p, PC-3p-73204_81, -3p-7719_599, and -3p-8403_561 were upregulated, and hsa-miR-190a-5p was down-regulated in older individuals (Fig. S5c; Table S6c), indicating that these miRNAs have a significant role in aging and could serve as potential markers of aging. Pathway analysis of the six significantly altered miRNAs revealed diverse pathways are associated with these miRNAs that could exacerbate underlying mechanisms of aging (Fig. S5d). While it is not entirely clear what the function of these miRNAs is it will remain of interest to monitor and report this important exosome cargo.

### Senescence and aging share mutual exosome signatures that may regulate age-related inflammation and ECM remodeling, potentially causing secondary senescence

We demonstrate a substantial overlap between the exosome proteins identified (52) and altered pathways between senescent fibroblasts and plasma from older individuals (Fig. 6B). Differential profiles of selected exosome proteins in aging and senescence, such as Antithrombin-III (SERPINC1), Prothrombin (F2), Coagulation factor V (F5), Plasminogen (PLG), Reelin (RELN), Soluble scavenger receptor cysteine-rich domain-containing protein SSC5D, Complement C1 subcomponents (C1r and C1s), and HLA class I histocompatibility antigen, A-24 alpha chain (HLA-A), showed complementary trends and were particularly associated with inflammation and ECM remodeling (Fig. 6B). Our DIA-MS proteomics analysis of senescent fibroblasts identified Plasminogen activator inhibitor 1 (SERPINE1), SPARC, Insulin-like growth factor-binding protein 7 (IGFBP7), Transforming growth factor beta-1 proprotein (TGFB1|1), and Integrin alpha-4 (ITGA4) as potential exosome protein candidates that can cause secondary senescence locally and distally (Fig. 6A). Our lipidomics data revealed the presence of lipids such as hexceramides HexCer(d18:1/24:0), HexCer(d18:1/24:1), and HexCer(d18:2/24:0), which induce secondary senescence [30], and lysophosphatidylcholine (LPC) class lipids that are involved in ECM modulation through metalloproteins. When comparing to the three different senescence-inducing stimuli on IMR90 cells, we saw similarly affected pathways with human plasma exosome aging. We observed a decrease in antioxidant and matrisome proteins, but more interestingly changes in inflammatory and ECM-modifier associated proteins (Fig. 7), that play crucial rolesin senescence. Dysregulation of the ECM composition, shape, firmness, and abundance contribute to pathological conditions, such as fibrosis and invasive cancer.

**Fig. 6.**
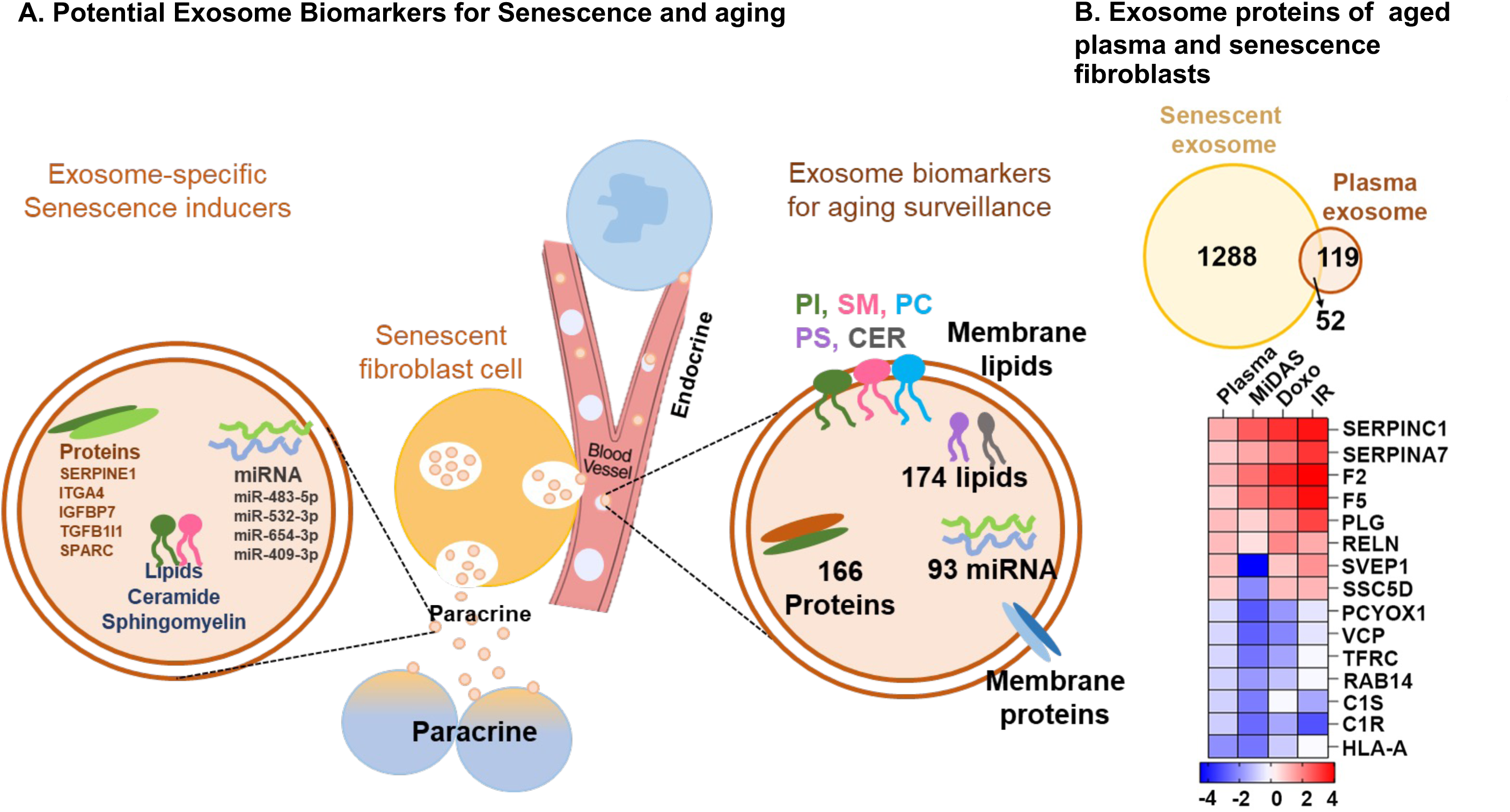
Overview of top potential exosome biomarkers for senescence and aging. **A.** Exosome proteins and miRNAs identified as potential biomarkers for secondary senescence (paracrine) and aging. **B.** Venn diagram showing overlapping and unique exosome proteins in senescent primary human lung fibroblasts (IMR90) and plasma from aging cohorts. Heatmap showing the top 15 commonly altered proteins in senescent fibroblasts and plasma from aging cohorts.

**Fig. 7.**
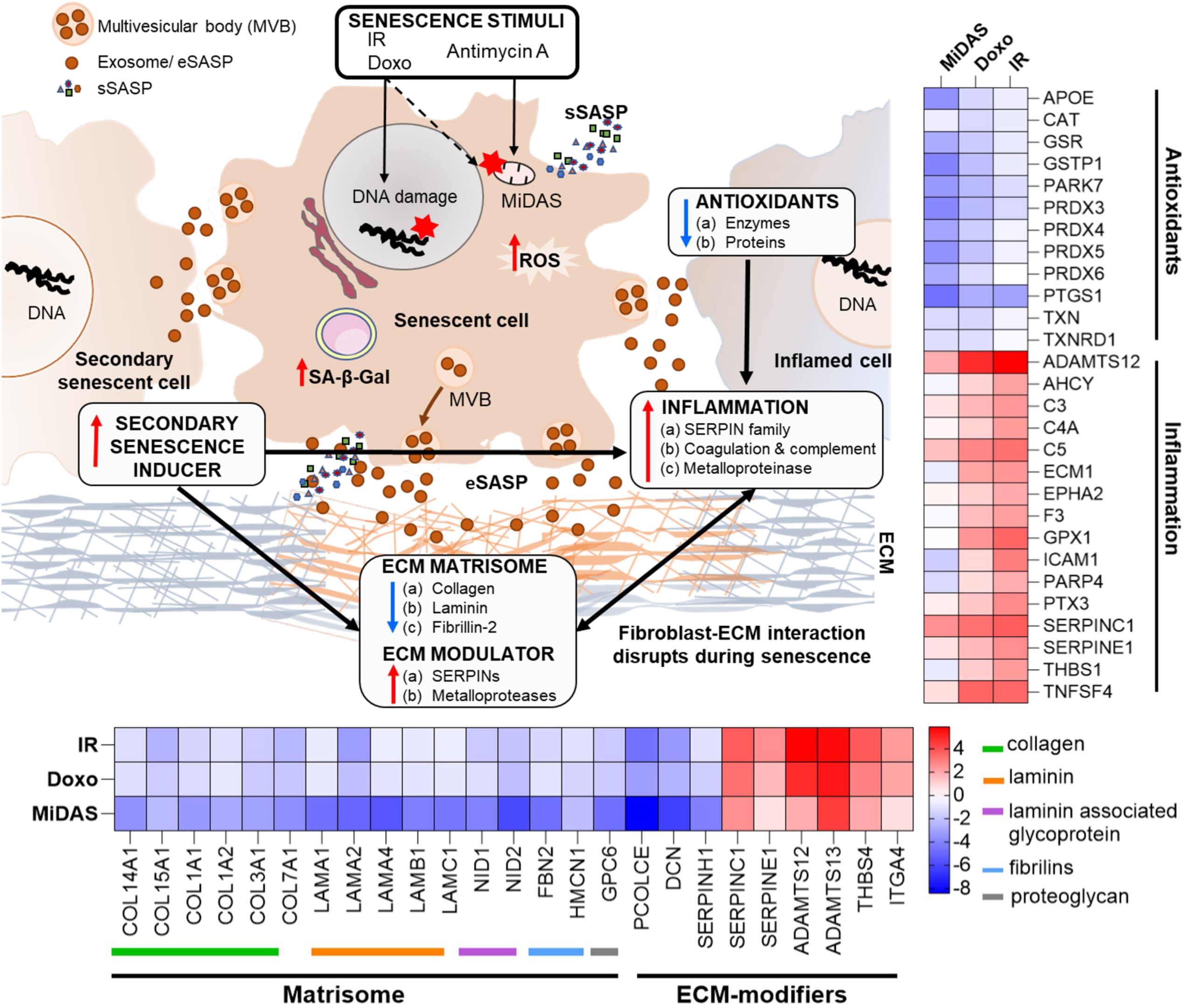
Summary of exosome-mediated intercellular communication in senescence and aging. A summary of our overall exosome proteomic findings from senescent human primary lung fibroblast cells that suggest potential mechanisms by which senescence is exacerbated and propagated to cause secondary senescence. Changes in inflammatory, antioxidants, and matrix remodeling-related proteins in exosomes (eSASP) from senescent cells may contribute to creating a senescent microenvironment that may influence and affect neighboring cells. Heatmaps show the top selected senescent fibroblast exosome proteins involved in inflammation, antioxidant, and ECM-remodeling pathways.

## Discussion

In recent years, several studies highlighted that the SASP secreted from senescent cells is highly complex and very heterogeneous depending on tissue type, cell type and origin of senescence or senescence induction [10, 31]. The composition of SASP can vary depending on a wide range of factors, including cell type, senescence-inducing stimuli, and age. While there has been significant progress in understanding SASP and its effect on neighboring cells and the extracellular matrix, what is currently known is just the tip of the iceberg. Exosomes have been recognized as a constituent of the SASP and are known to play a crucial role in facilitating intercellular communication. Exosomes secreted from senescent cells are still relatively underexplored. Understanding the complexity, cell-, tissue-, and stimuli-specificity of exosomes from senescent cells could offer essential insights into the mechanisms underlying the progression of senescence, aging, and age-related diseases. These insights could lead to the development of potential diagnostic and therapeutic interventions.

We have laid out a comprehensive and quantitative DIA-MS proteomic workflow for exploring exosome profiles from different senescence-inducing stimuli in cells and plasma. Isolation and enrichment of intact and contamination-free exosomes are challenging, but we propose that exosomes isolated by sequential SEC and UF improve the accuracy and reliability of downstream proteomics analysis. This is based on specific sizes and surface markers that distinguish exosomes from other extracellular vesicles and low levels of protein contaminants, which can interfere with downstream analysis (Fig. 2). This approach led to the identification of 60-70% exosome-specific proteins in IMR90 fibroblasts (Fig. 3) and plasma (Fig. 4).

The limited availability of high-quality exosome libraries is another challenge. Spectral library searching is a prerequisite for DIA quantitative proteomic analysis. We present three high-quality human plasma exosome spectral libraries (DDA, directDIA, and hybrid), cataloging around 2,700 high-confidence proteins as a resource for exosome proteomic research (Fig. S2). The hybrid library identified a more considerable amount of total and significantly altered proteins, while the directDIA approach is instrumental in cases where no pre-existing large libraries are available, such as in non-model organisms or tissues. We emphasize the significance of <u>comprehensive quantitative workflows using DIA-MS proteomics for exosome</u> <u>research</u>. These mass spectrometric approaches offer highly robust workflows for biomarker discovery in clinical samples, and specifically for biofluids and circulating biomarkers.

Previously we reported the profiling of soluble proteins and exosomes from senescent cells and their SASP which led to the creation of the SASP Atlas [10]. Here we build on our prior research by leveraging the DIA exosome proteomics workflow. This novel approach allowed us to detect changes in the exosome protein profiles between young and old individuals in human plasma, as well as from senescent IMR90 fibroblasts induced by IR, doxo, or MiDAS (Fig. 3). Studying senescence and SASP with different senescent-inducing methods is essential to gain a more comprehensive understanding of the SASP and identify commonalities and differences in the composition, which can illuminate how the SASP components propagate and exacerbate the senescent phenotype.

Our examination of SASP protein profiles in plasma samples from older individuals has provided us with a more comprehensive understanding of the changes in intercellular communication that occur with aging. This analysis encompasses exosomes derived from all cell types, tissues, and senescence stimuli. In our study, we have identified 1,288 and 119 exosome-specific proteins unique to senescent fibroblasts and plasma samples from aging cohorts respectively, while 52 proteins are common to both (Fig. 6B). By cross-analyzing exosomes derived from plasma and cells, we could begin to dissect their endocrine and paracrine mode-of-action in the context of senescence [4], which might potentially play a role in secondary senescence. Johnson *et al.* (2020) conducted a meta-analysis of 36 proteomics studies, identifying 818 aging-associated soluble proteins in the plasma proteome that were altered in at least two studies [32]. Of these, 28 proteins overlapped with our age-associated plasma exosome panel, 228 proteins overlapped with our exosome SASP components, and notably, 14 proteins were common across all three datasets, as shown in Fig. S6 and Table S7. The limited overlap in proteins can be attributed to two main factors: first, we are comparing exosome SASPs and age-associated plasma exosome proteins with age-related soluble plasma proteins; second, our data were generated using mass spectrometry, whereas the Johnson *et al.* data were compiled from 36 studies using diverse proteomics methods.

Interestingly, several variants of immunoglobulins (n=61, as presented in Table S4c) show a strong negative correlation with aging. It suggests that the humoral immune response is downregulated in older individuals, which may contribute to the increased susceptibility to infections and diseases associated with aging (Fig. 4G). The downregulation of immunoglobulins in plasma exosomes from older individuals is consistent with previous studies that have reported age-related changes in the immune system, including a decline in B cell function and decreased antibody production. In contrast, SASP sustains and enhances inflammation, which is a broad immune response to tissue injury, infection, or other stimuli. While inflammation can be beneficial in response to acute infections or injuries, chronic inflammation can contribute to various age-related diseases. For example, exosomes released by prostate cancer cells carry pro-inflammatory M2 phenotype and promote a pro-tumorigenic microenvironment [33]. Here we demonstrate that the specific inflammatory proteins that are upregulated in the exosome SASP can vary depending on the senescence-inducing stimulus, but some common examples are serine protease inhibitors (SERPIN family, SERPINC1), Prothrombin (F2), Coagulation Factor V (F5), Reelin (RELN), Disintegrin and metalloproteinase with thrombospondin motifs 12 (ADAMTS12), Thrombospondin-1 (THBS1), Complement C3, and Complement C5 (Fig. 6-7). Further, we observed decreased levels of antioxidant proteins in senescent exosomes (Fig. 7). Biochemical analysis reveals an overall increase in intracellular oxidative stress in all senescent conditions, the highest being in MiDAS (Fig. 3J). Chronic oxidative stress and inflammation are closely linked and can lead to the development and progression of many age-related diseases, such as cancer, cardiovascular diseases, and neurodegenerative disorders [34]. In addition, inflammatory signals can degrade ECM components and alter the ECM structure [35].

Changes in the extracellular matrix (ECM) play a crucial role in senescence and cancer, as senescent cells alter the ECM by secreting ECM remodeling components. However, the molecular understanding of the interplay between ECM and senescent cells is limited. Our study revealed that exosomes secreted by senescent cells exhibit decreased level of ECM structural proteins, including collagens, laminins, and Fibrillin-2 (Fig. 7). This suggests a potential mechanism by which senescent cells destabilize the ECM, contributing to the propagation of a senescent microenvironment. In addition, a decreased abundance in collagen mRNA levels is noted in senescent retinal pigment epithelial cells, which are major ECM producers in the retina, and hematopoietic stem cells [36]. ECM alterations are widely recognized as a significant factor driving abnormal cellular responses [37], and recent research indicates that fragmentation of collagen fibrils during aging leads to the disruption of fibroblast-ECM interactions [38]. This disruption can contribute to the development of an “aged” phenotype. Our findings and those of others strongly suggest that ECM and senescence play a critical role in aging.

Inflammation and ECM modulation might be two key factors contributing to secondary senescence, leading to the accumulation of senescent cells and further promoting inflammation and ECM remodeling. Recent studies suggest that sEVs can transfer senescent factors to non-senescent/healthy cells [13, 39]. We identified multiple proteins, lipids, and miRNAs relevant to paracrine senescence signaling that are upregulated in exosomes released by senescent fibroblasts. Notable secondary signals include integrin alpha-4 [40], SERPINE1 [41, 42], insulin-like growth factor-binding protein 7 [43, 44], transforming growth factor beta-1 proprotein [45], SPARC [46], and hexceramides (HexCer) [30]. Additionally, miRNAs such as miR-483-5p [26], miR-532-3p [26], and miR-409-3p [47] were consistently elevated. Our studies offer valuable insights into the potential role of exosomes in SASP-mediated paracrine senescence, offering potential therapeutic targets for mitigating its effects.

Our lipidomic analysis revealed consistent upregulation of phospholipid classes, including phosphatidylcholine (PC) and phosphatidylethanolamine (PE), aligning with previous studies in senescent cells [48]. Furthermore, we observed significant increases in PS, SM, and ether-linked/plasmalogen PE species (PE(O/P-)) across all senescence inducers, implicating altered exosomal lipid composition in senescence-associated changes in cellular functions, particularly membrane dynamics. Conversely, lower levels of specific triglycerides (TG) and acylcarnitines (AC) were noted, except in MiDAS-induced senescence, possibly reflecting reduced TG accumulation [49] and decreased AC synthesis during senescence. Finally, while 112 lipids were commonly altered across all senescence inducers, distinct lipid signatures were observed for specific stimuli, such as doxo (7 unique lipids) and MiDAS (9 unique lipids; Table S5c), highlighting stimulus-specific molecular mechanisms of senescence. These findings deepen our understanding of exosome-mediated signaling in senescence and point toward tailored approaches for therapeutic intervention.

Increasing evidence of age-associated changes in miRNA levels is seen in various models, ranging from nematodes to humans [38]. We reproducibly identified 88 unique miRNAs, including miR-27a-5p, -874-3p, in plasma exosomes from older individuals. An upregulation of miR-27a is reported in degenerative diseases such as osteoarthritis [50] and AD [51]. Moreover, miR-874 was identified in circulating brain fluid from patients with mild cognitive impairment [52] and as a biomarker for early AD development [52, 53]. These miRNAs could be developed into future senescence biomarkers. Further investigations within larger aging populations and longitudinal analysis are needed to confirm the potential of these protein, lipid, miRNA candidate signatures as diagnostic and prognostic biomarkers.

Overall, we describe an exosome DIA-proteomic and multi-omic workflow for exploring exosome profiles from cells and plasma. We emphasize the power and potential of using DIA-MS proteomic data analysis for exosome research as a highly robust workflow for biomarker discovery in clinical samples. Our study reveals that the exosome SASPs exhibit considerable heterogeneity, and we systematically report the changes in exosome cargo, specifically proteins, lipids, and miRNAs, between young and old individuals and senescent fibroblasts induced by IR, doxo, or MiDAS. We also report that exosomes may be critical players in mitigating senescence and aging by inflammation, oxidative stress, ECM modulation, and secondary senescence. Additional investigations and functional validations are required to ascertain the mechanisms by which exosomes mediate their effects in the context of senescence and aging and understand how the senescence- and age-associated exosome cargos can impact the recipient cells. This can help understand the global molecular mechanisms underlying aging and age-related diseases and potentially identify novel therapeutic targets or biomarkers to target and regulate senescent cell-secreted exosomes or use them as therapeutic agents.

## Methods

### Human Plasma Samples

We obtained 10 plasma samples, 5 from young individuals (20-26 years old) and 5 from older individuals (65-74 years old), with no restrictions regarding gender, from the Blood Centers of the Pacific in San Francisco, CA, detailed in Table S1. Specimens from individuals with any known autoimmune disorders, chronic diseases, bleeding disorders, or neurodegenerative disorders were excluded. The plasma samples were aliquoted and stored at -80°C until further use. IRB approvals were not required for the plasma samples.

### Primary human fibroblast tissue culture and senescence induction

#### (a) Primary human fibroblast tissue culture

Primary human lung fibroblasts (IMR-90) used in the study were acquired from ATCC, Manassas, VA (#CCL-186). Cells were maintained in Dulbecco’s Modified Eagle Medium (DMEM) supplemented with penicillin and streptomycin (5,000 U/mL and 5,000 μg/mL; Thermo Fisher Scientific, Waltham, MA, #15070063) and 10% fetal bovine serum (FBS; Thermo Fisher Scientific, Waltham, MA, #2614079), and incubated at 37°C in an atmosphere of 10% CO_2_ and 3% O_2_.

#### (b) Senescence induction in primary human fibroblasts

Irradiation (IR), doxorubicin (doxo), and Antimycin A induced-mitochondrial dysfunction-associated senescence (MiDAS) were used to induce senescence in IMR90 cells. For IR induction, the cells were irradiated with ionizing radiation (10 Gy) for 30 minutes. For doxo induction, the cells were treated with 350 nM doxo (Thermo Scientific #25316-40-9) for 24 h. For MiDAS induction, 200 μM Antimycin A (AA; Thermo Scientific # 1397-94-0) was supplemented to the complete media every alternate day for 10 d. After senescence induction, the cells were cultured for 10 d to allow the senescent phenotype to develop. The conditioned media was collected for exosome analysis after the cells were washed with phosphate-buffered saline (PBS; Gibco™ #10010023) and placed in serum- and phenol-red-free DMEM media for 24 h. In parallel, the control cells were made quiescent by incubating in 0.2% serum (minimal media) for 3 d by mock irradiation or treated with DMSO in 0.2% serum for 1 d. Subsequently, senescence-induced and control fibroblasts were placed in serum- and phenol-red-free DMEM media for 24 h before collecting the conditioned media for exosome analysis.

#### (c) Cell viability assays

On day 10 for senescence-inducers or day 3 for control quiescent cells (as shown in Fig. 3A (ii)), cell viability was assessed using SYTOX Green (Invitrogen, Carlsbad, CA, #S7020). The protocol recommended by the manufacturer was followed, which involved staining the cells with SYTOX, and counting the nuclei that exhibited staining to indicate cell death using a fluorescent microscope (Evident, #Olympus IX70 Fluorescence) with a wavelength of 523 nm.

#### (d) EdU and SA-β-Gal staining

The activity of SA-β-gal (senescence-associated β-galactosidase) was determined using a BioVision Senescence Detection Kit (Milpitas, CA, #K320-250), which is specifically designed to measure the activity of lysosomal β-galactosidase enzyme that is known to increase significantly in senescent cells. To perform the assay, 7000 cells for each condition, including replicates, were seeded into 8-well culture slides coated with poly-lysine (Corning, Corning, NY; #354632). After 24 h, the cells were stained with the SA-β-gal detection reagent according to the manufacturer’s protocol. In addition to measuring SA-β-gal, we evaluated cell proliferation using the Click-iT EdU Cell Proliferation Assay Kit (Invitrogen, #C10337). This kit utilizes a thymidine analog, EdU, which is incorporated into newly synthesized DNA during the S-phase of the cell cycle. The cells were treated with 10 μM EdU for 24 h, then fixed and permeabilized. The incorporated EdU was detected using a Click-iT reaction cocktail that binds to the EdU and produces a fluorescent signal. The EdU and SA-β-Gal stained cells were then imaged using a Fluorescence microscope (Olympus USA, # IX70) and quantified using Image J software (IJ 1.46r, NIH). Overall, EdU and SA-β-Gal assays allow for measuring both senescence and cell proliferation in response to different senescence-induction.

#### (c) NAD+/NADH ratio determinations

We measured NAD+ and NADH levels of Antimycin A-treated MiDAS-induced senescent, and Qui control IMR90 cells using a commercial colorimetric NAD/NADH Assay Kit (Abcam, UK, #ab65348). The assays were performed in biological triplicate, with 2 × 10^6^ IMR90 cells lysed in NAD/NADH extraction buffer by two freeze/thaw cycles (20 minutes on dry ice followed by 10 minutes at RT), followed by centrifugation (14,000 X g for 5 min at 4°C) and filtration (10 kDa cut-off filters, MilliporeSigma™, #UFC501024) to remove cellular debris and proteins. The resulting supernatant was then subjected to a series of enzymatic reactions that converted NADH to a fluorescent product (absorbance at 450 nm) as per the manufacturer’s instructions, allowing for quantitative measurement of the NAD+/NADH ratio.

#### (d) Oxidative stress measurements

Lipid peroxidation is a common consequence of oxidative stress. A lipid peroxidation assay kit (Abcam, UK, #ab233471) was used to measure cellular oxidative stress in IR-, doxo-, and MiDAS-induced senescent fibroblasts. Qui or DMSO fibroblasts were used as a control for comparison. Briefly, 5 × 10^5^ cells in triplicates for each condition were homogenized in 500 μL of 20mM Na Phosphate buffer (pH about 3 to 3.2) and 0.5% TritonX-100 to solubilize the cell membranes, followed by centrifugation (14,000 X g for 5 min at 4°C) to remove cellular debris. The resulting supernatant was then subjected to a series of chemical reactions per the manufacturer’s instructions to produce a colored product, which was quantified spectrophotometrically at 695 nm.

### Exosome isolation and enrichment

#### (a) Human plasma

Ultracentrifugation (UC), sequential SEC coupled with either ultrafiltration (SEC/UF) or ultracentrifugation (SEC/UC) was used to isolate and enrich exosomes from plasma. Plasma was centrifuged at 3,000 x g for 10 min to remove cellular debris and then centrifuged at 10,000 x g for 10 min to remove apoptotic cell bodies. The supernatant (2 mL) containing extracellular vesicles was then proceeded by UC, SEC/UF, or SEC/UC.

For SEC, the supernatant was passed through a qEV size-exclusion column (Izon, #qEV35) for exosome enrichment, according to the manufacturer’s instructions. Briefly, the column was maintained at room temperature for 20 min before use and then activated by flushing with 1.5x column volume of degassed PBS (∼60 mL). The supernatant was passed through the activated qEV size exclusion columns (by gravity drip). Different fractions were collected by filling the reservoir with PBS (∼50 mL). The initial 14 mL of flow-through was void volume, while the subsequent 10 mL (collected in 5 x 2 mL centrifuge tubes: labeled as E1–5) was enriched exosome fractions based on immuno-blotting with exosome-specific protein markers (Fig. 2). The remaining 26 mL was marked as the post-exosome fractions that contained plasma proteins. qEV size-exclusion exosome fraction (E1–5) were quality-checked spectrophotometrically (SpectraMax Plus 384, Molecular devices), where OD600 was used for exosome quantification and OD280 for plasma proteins. Based on higher plasma protein contamination in fraction E5, we pooled only fractions E1–E4 to obtain highly pure exosomes. Subsequently, fractions E1–E4 were concentrated by ultrafiltration (UF; Amicon® Ultra 50 kDa, # UFC903024) or ultracentrifugation (Optima, The Beckman Coulter Life Sciences) for further exosome analysis.

#### (b) DMEM-conditioned media

50 mL of conditioned media from IR, doxo, MiDAS-senescent, and Qui-control IMR90 cells were centrifuged at 3,000 x g for 10 min to remove any cell debris. The supernatant was passed through a 50 kDa cut-off filter. PBS was added to the remaining filtrate to make up to 2 mL. The filtrate was then centrifuged at 15,000 x g for 30 min to remove apoptotic cell bodies and proceeded with either SEC/UF or Antibody-based extractions (AbE). For SEC/UF, the supernatant was passed through a size exclusion chromatography column followed by ultrafiltration to enrich and purify the exosomes, as described previously.

For AbE enrichment a Miltenyi Biotec exosome isolation kit (Miltenyi Biotec, #130-110-912) was used to label the exosomes with magnetic microbeads that specifically bind to exosome surface markers. Briefly, 2 mL of filtrate was added to 50 µL of magnetic microbeads and incubated for 1 h at room temperature on a rotor. A micro-column was equilibrated with 100 µL of equilibration buffer and washed thrice with 100 µL of isolation buffer. The labeled exosomes were then passed through a micro-column, and the exosomes were selectively retained on the column while other non-exosome particles were removed. Finally, the exosomes were washed 4x with 100 µL isolation buffer and eluted with 200 µL of elution buffer. We used SEC/UF for proteomics analysis, while AbE was used for lipidomics studies.

### Exosome quality check

#### (a) Size distribution analysis

Exosome diameter was assessed by tunable resistive pulse sensing (TRPS) on an IZON qNano Gold nanoparticle characterization instrument using an NP150 nanopore membrane at calibration with 110-nm carboxylated polystyrene beads at a concentration of 1 × 10^10^ particles/mL (Zen-bio, Research Triangle, NC).

#### (b) Immunoblotting

Exosomes were lysed in RIPA buffer (Thermo Scientific™, #89900) for protein extraction. Exosome proteins were quantified using BCA (Thermo Scientific™, #23225). 20 μg of protein/well was separated on 12% SDS-PAGE and transferred onto PVDF membranes under semi-dry conditions using the Trans-Blot Turbo Transfer System transfer unit (Bio-Rad, #1704150). Ponceau S staining (Sigma-Aldrich, #P7170-1L) confirmed equal loading of exosomal proteins. Immunoblotting was performed using 1:1000 dilution of CD9 (Abcam, #Ab96696), Tumor susceptibility gene 101 (TSG101, Santa Cruz Biotechnology, #sc-7964), albumin (Abcam, #Ab106582), or Immunoglobulin G (IgG, Abcam, #Ab200699), and anti-rabbit HRP-conjugated antibody (Bio-Rad, #1706515) or anti-mouse HRP-conjugated antibody (Abcam, #Ab6728) as a secondary antibody (1:3000 dilution). Blots were developed per the manufacturer’s protocol with the Pierce ECL Western Blotting Substrate (Thermo Fisher Scientific, #32106). Imaging was performed by Azure Biosystems c600. Densitometric analysis of bands was performed using Image J software (IJ 1.46r; NIH).

### Mass spectrometric analysis of exosome proteins

#### (a) Protein extraction and trypsin digestion

Plasma exosomes were lysed in lysis buffer (8 M urea, 0.1 M ammonium bicarbonate), and then reduced by 10 mM tris(2-carboxyethyl) phosphine (TCEP) and alkylated using 40 mM chloroacetamide (CAA) for 1 h at 37°C. The solution was diluted with 0.1 M ammonium bicarbonate buffer to lower the urea concentration to 1.5 M and digested with trypsin (Promega, Madison, WI, # V5111) at a 1:100 ratio (protease to protein (wt/wt)) overnight at 37°C. Peptides were purified using MacroSpin clean-up columns (NEST group, Southborough, MA, # SEM SS18V), following the manufacturer’s protocol. Eluted peptides were vacuum dried and then resuspended in 60 µL of 1% acetonitrile acidified with 0.1% formic acid buffer spiked with iRT peptides (Biognosys, Schlieren, Switzerland, # Ki-3002-1). Peptides were quantified using nano-drop (Spectrostar Nano, BMG Labtech, # UV5Nano) and adjusted to a 1 mg/mL concentration.

IMR90 cell exosomes were lysed in the lysis buffer (5% SDS and 50 mM TEAB), reduced, alkylated, and acidified to a final concentration of 1.2% phosphoric acid. Then 90% methanol in 100 mM TEAB was added and passed through micro S-Trap columns (Protifi, #C02-micro-80). IMR90 exosome proteins trapped in the S-Trap columns were digested with trypsin (Promega, Madison, WI, # V5111) in digestion buffer (50 mM TEAB, pH ∼8) at a 1:25 ratio (protease to protein (wt/wt)) overnight. Peptides were eluted from the S-Trap column with 50 mM TEAB and 0.5% formic acid and then with 50% acetonitrile in 0.5% formic acid. Pooled elution solutions were dried in a speed vac and resuspended in 0.2% formic acid. The resuspended peptide samples were desalted, concentrated, and resuspended in aqueous 0.2% formic acid containing ‘Hyper Reaction Monitoring’ indexed retention time peptide standards (iRT, Biognosys, #Ki-3002-1).

#### (b) Reversed-phase fractionation (RP) for deep spectral library generation

For DDA library generation, 400 µg of plasma exosome peptides (pooled from 10 different plasma exosome samples) were injected onto Dionex Ultimate 3000 LC (Thermo Scientific) coupled to ACQUITY UPLC CSH1.7 mm C_18_ column (2.1 x 150 mm) (Waters, Milford) for fractionation using RP. Peptides were chromatographically separated in a 30-min non-linear gradient from 1% to 40% RP buffers (Buffer B: 100% ACN; Buffer A: 20 mM ammonium formate, pH 10). A micro fraction was taken every 45 seconds, pooled into 24 final peptide fractions, vacuum dried, resuspended in 20 mL buffer A spiked with iRT peptides, and quantified. Each fraction was acquired in DDA mode, as described below.

#### (c) LC-HCD-MS-based exosome proteins data acquisition

2 µg of trypsin-digested plasma exosome peptides were separated using liquid chromatography coupled to a mass spectrometer. The analytical column was in-house packed into a fritted tip emitter to a length of 50 cm (ID 75 µm) (New Objective, Woburn, MA) using the CSH C_18_ phase (1.7 µm) (Waters, Milford, MA). The column was operated using an Easy nLC 1200 (Thermo Fisher Scientific, San Jose, CA) coupled online to an Exploris 480 spectrometer (Thermo Fisher Scientific). Peptides were eluted at 250 mL min-1 using a non-linear 2-h gradient from 1% to 45% buffer B (Buffer B: 80% ACN + 0.1% FA; Buffer A: 0.1% FA). Based on the RP samples, the mass spectrometer was operated in a data-dependent mode for DDA library generation. Briefly, the following settings were applied: MS1 scan resolution: 60,000; MS1 AGC target: 300; MS1 maximum IT: 25 ms; MS1 scan range: 350–1650 Th; MS2 scan resolution: 15,000; MS2 AGC target: 200; MS2 maximum IT: 25 ms; isolation window: 4 Th; first fixed mass: 200 Th; NCE: 27; minimum AGC target: 1e3; only charge states 2 to 6 considered; peptide match: preferred; dynamic exclusion time: 30 s.

In DIA mode, all samples were acquired in a randomized fashion with regard to the different conditions: young and old. The mass spectrometer was operated using the following parameters for the MS1 scan: scan range: 350 to 1650 Th; AGC target: 300%; max injection time: 20 ms; scan resolution: 120,000. The MS1 was followed by targeted MS2 scan events with the following settings: AGC target: 1000%; max injection time: 54 ms; scan resolution: 30,000; scan range: 350-1,650 m/z Th; normalized collision energy: 27. The number of DIA segments and the segment widths were adjusted to the precursor density and to achieve optimal data points across each peak for each acquisition (Table S8a). The DIA methods consisted of 40 DIA segments with a cycle time of 3.2 s.

IMR-90 exosome peptides were separated on a Dionex UltiMate 3000 system, and DIA-MS acquisition was performed on an Orbitrap Eclipse Tribrid mass spectrometer (Thermo Fisher Scientific, San Jose, CA). The solvent system consisted of 2% ACN, 0.1% FA in H2O (solvent A), and 98% ACN, 0.1% FA in H2O (solvent B). Proteolytic peptides (50 ng) were loaded onto an Acclaim PepMap 100 C_18_ trap column (0.1 x 20 mm, 5 µm particle size; Thermo Fisher Scientific) for 5 min at 5 µL/min with 100% solvent A. Peptides were eluted on an Acclaim PepMap 100 C_18_ analytical column (75 µm x 50 cm, 3 µm particle size; Thermo Fisher Scientific) at 300 nL/min using the following gradient of solvent B: 2% for 5 min, linear from 2% to 20% in 125 min, linear from 20% to 32% in 40 min, up to 80% in 1 min, 80% for 9 min, down to 2% in 1 min, and 2% for 29 min, for a total gradient length of 210 min. Full MS spectra were collected at 120,000 resolution (AGC target: 3e6 ions, maximum injection time: 60 ms, 350-1,650 m/z), and MS2 spectra at 30,000 resolution (AGC target: 3e6 ions, maximum injection time: Auto, NCE: 27, fixed first mass 200 m/z). The isolation scheme consisted of 26 variable windows covering the 350-1,650 m/z range with an overlap of 1 m/z (Table S8b) [54].

#### (d) Exosome protein spectral library generation from DDA and DIA

Three spectral libraries were generated i) a deep DDA-based spectral library, ii) a DIA-only spectral library (directDIA), and iii) a hybrid spectral library (hybrid) (Fig. 1). The DDA deep spectral library was generated by performing DDA analysis on 25 RP fractions representing fractions from a pooled mixture of peptides obtained from the same 10 plasma exosome protein samples (n = 5, young and old each). DDA raw files (n = 25) were individually searched with Pulsar in Spectronaut version 14.0.200601.47784 (Biognosys) against the Human UniProt FASTA (downloaded on January 31, 2018, containing 92,931 proteins) using the following settings: fixed modification: carbamidomethyl (C); variable modifications: acetyl (protein N term), oxidation (M); enzyme: trypsin/P with up to two missed cleavages. Spectronaut automatically determined mass tolerances and other settings were set to default. Search results were filtered using a 1% false discovery rate on the precursor ion, peptide, and protein levels [55, 56]. For the directDIA spectral library, 10 individual DIA acquisitions of plasma exosomes (n = 5, young and old each) were processed using Pulsar in Spectronaut version 14.0.200601.47784 using the same human FASTA file and settings as above. We combined both DDA and directDIA libraries for the hybrid library generation in Spectronaut. All Spectronaut parameter settings were uploaded to the data repository.

#### (e) DIA data processing and quantification

Quantitative analysis was performed by processing protein peak areas determined by the Spectronaut software. Prior to library-based analysis of the DIA data, the DIA raw files were converted into htrms files using the htrms converter (Biognosys). MS1 and MS2 data were centroided during conversion, and the other parameters were set to default. For the human plasma exosome cohort, the htrms files were analyzed with Spectronaut (version: 14.0.200601.47784, Biognosys) using the previously generated libraries to perform quantitative data analysis with the three libraries generated (see above): the directDIA library, the deep DDA spectral library, and the hybrid library. Briefly, calibration was set to non-linear iRT calibration with precision iRT selected. DIA data was matched against the described spectral library supplemented with decoys (library size fraction of 0.1) using dynamic mass tolerances. Quantification was based on MS/MS XICs of 3-6 MS/MS fragment ions, typically y- and b-ions, matching specific peptides in the spectral library. Interference correction was enabled on MS1 and MS2 levels. Precursor and protein identifications were filtered to 1% FDR, estimated using the mProphet algorithm, and iRT profiling was enabled. Quantification was normalized to the local total ion chromatogram. Exosome proteins identified with less than two unique peptides were excluded from the analysis. The DIA data was processed for relative quantification comparing peptide peak areas from different conditions (old vs. young plasma and senescent vs. Qui control IMR90 fibroblasts). Differential analysis was performed applying paired t-test, and p-values were corrected for multiple testing. Protein groups with q-value ≤ 0.05 and log_2_ ratio ≥ 0.58 were considered significant. For the IMR90 exosome cohort, DIA data were analyzed with Spectronaut (version: 15.1.210713.50606, Biognosys) without a spectral library using the directDIA algorithm. Similar settings were applied for DIA data processing and differential analysis.

### Mass spectrometric exosome lipid analysis

#### (a) Exosome lipid extraction

AbE-based enriched exosome lipids were extracted by a modified Folch extraction method [57, 58]. Briefly, 200 µL of fibroblast (IMR-90) exosomes in methanol were also transferred into a 1.7 mL Sorenson microcentrifuge tube (# 89082-332) for lipid extraction. The IMR90 cell exosome count ranged from 10-13 million and was used for IMR90 sample normalization. Each IMR90 exosome sample (200 µL) was initially diluted in 200 µL in Optima LC-MS grade methanol, where the IMR90 cell exosomes were normalized to 10 million exosomes present. Each sample was dried in a Thermo Scientific Speedvac SPD130DLX (San Jose, CA) for 30 min, and then 750 µL of chilled methanol was added to each tube. The samples were then transferred to a 2.0 mL bead tube that contained 2.4 mm tungsten carbide beads (Fisher Scientific, #15-340-153) before being placed into a Fisher Brand Bead Mill 24 Homogenizer (#15-340-163) for 2 min at a frequency of 30 Hz for 2 cycles. Each sample was then placed into a 5.0 mL glass vial (Fisher Scientific, #S28023) equipped with a Teflon-lined cap, and 750 µL of methanol was added. Next, 3.0 mL of chloroform (Sigma Aldrich LC-MS grade) and 200 µL of ultrapure water (Optima LC-MS grade) were added to induce phase separation. Samples were then vortexed with a VWR Standard Heavy-Duty Vortex Mixer for 30 s before being placed in a Branson CPX5800H sonicator for 30 min at room temperature. Once completed, the samples were incubated at 4°C for 1 h to achieve the phase separation. For each sample, 1.20 mL of water was added, and mixed gently. Next, the samples were placed in an Eppendorf 5810R Centrifuge and centrifuged at 1000 rpm for 10 min. Finally, 300 µL of the bottom organic layer of each sample was collected and dried in a speed vac for approximately 40 min until dry. The total lipid extracts were finally reconstituted with 190 µL of methanol and 10 µL of chloroform and stored at -20°C until mass spectrometric analyses were conducted the following week. Before mass spectrometric measurements, stored samples were dried and reconstituted in the 190 µL:10 µL methanol:chloroform solvent mixture.

#### (b) LC-IMS-CID-MS-based exosome lipids data acquisition

An Agilent 1290 Infinity II UHPLC (Santa Clara, CA) coupled with an Agilent 6560 IM-QTOF mass spectrometer (Santa Clara, CA) was used for the LC-IMS-CID-MS analysis of IMR90 exosome lipids [59]. For the exosome lipid acquisition, 10 µL of extracted exosome lipids were injected onto a reversed-phase Waters CSH column (3.0 mm x 150 mm x 1.7 µm particle size) and chromatographically separated over a 34 min gradient period (Mobile Phase A: 10 mM NH_4_AC in ACN/H_2_O (40:60); Mobile Phase B: 10 mM NH_4_ ACN/IPA (10:90)) at a flow rate of 250 µL/min [60]. Details regarding the gradient and column wash are provided in Table S8c. Lipids were subsequently analyzed using positive and negative ionization modes using an Agilent Jet Stream ESI source (Agilent Technologies, Santa Clara, CA). The lipids were focused, trapped, and pulsed into the IMS drift cell using a high-pressure and trapping ion funnel with pressures ∼4 torrs [59]. Ions were then separated with drift tube IMS (DTIMS) in the 78 cm drift tube following the ion funnel. Ions were then transferred to a hexapole collision cell where alternating precursor and all-ions fragmentation scans were acquired. Collision energies for the collision-induced dissociation (CID) analyses were ramped based on the various IMS drift times for the different lipid species of interest, providing optimized fragmentation for the varying ion sizes as well as 1+ and 1- charge states [61]. The collision energy ramp applied based on IMS drift time is shown in Table S8d. Finally, various ion optics transferred the ions to the time-of-flight (TOF) mass spectrometer. The TOF mass spectrometer was set to evaluate a mass range of 50 to 1700 *m/z,* and the LC, IMS, and MS information were collected in an Agilent .d file. Brain total lipid extract (BTLE; Avanti Polar Lipids, Alabaster, AL #131101C) was used as quality control (QC) for the LC, IMS, and MS values and observed peak abundances.

#### (c) Lipid Identification

For lipid identification, we considered LC retention times, IMS collisional cross section (CCS), and m/z values of the precursor and fragment ions with Skyline software and a lipidomic plasma library comprising 516 unique lipids published previously [62]. These lipids were all experimentally validated with LC, IMS, and MS information. Speciation of the lipids included the headgroup and fatty acyl (FA) components (e.g., PE (16:0_18:0)) but did not account for double bond position and orientation or specific backbone position. A total of 247 exosome lipids were identified in all samples (Table S5a).

For the lipidomic comparative analysis, the lipid intensities were total ion chromatogram (TIC) normalized and log_2_ transformed. To assess clustering, PLS-DA was performed using the package ‘mixOmics’. Significant lipids were identified in R using the package ‘limma’, which employs linear models to assess differential expression in complex experiments. The resulting p-values were adjusted using Benjamini-Hochberg correction and fold changes were log_2_ transformed. A heatmap of significant lipids was created using the package ‘pheatmap’ and group clustering was assessed using Euclidean distance and complete linkage. Finally, violin plots were created using the package ‘ggplot2’. The exported statistical outputs, including log_2_ fold changes and p-values for the IR vs. LSM (Qui), doxo vs. DMSO, and MiDAS vs. DMSO comparisons are provided in Tables S5b and S5c.

### Plasma exosome miRNA extraction, deep sequencing, and analysis

The SEC/UF extracted plasma exosomes were outsourced to LC Sciences, LLC (Houston, TX) for miRNA deep sequencing and analysis. Briefly, a small RNA (sRNA) library was generated for 10 plasma exosome samples (n=5, young and old individuals each) using a propriety Illumina Truseq™ Small RNA Preparation kit (Illumina, # RS-930), according to the manufacturers’ guide. The purified cDNA library was used for cluster generation on Illumina’s Cluster Station and then sequenced on Illumina GAIIx following the vendor’s instruction for operating the instrument. Raw sequencing reads (40 nts) was obtained using Illumina’s Sequencing Control Studio software version 2.8 (SCS v2.8), following real-time sequencing image analysis and base-calling by Illumina’s Real-Time Analysis version 1.8.70 (RTA v1.8.70). The extracted sequencing reads were used in the standard data analysis. A proprietary pipeline script, ACGT101-miR v4.2 (LC Sciences), was used for sequencing data analysis [63]. For the miRNA differential analysis, a p-value < 0.05, an absolute fold change > 1.5, and a read count > 0 in each condition were required to identify statistically-significant changes.

### Data availability

#### a. Exosome protein cargo

All mass spectrometry raw files, spectral libraries, Spectronaut files, descriptive methods, and other supplementary tables and data have been deposited to MassIVE ftp://MSV000086782@massive.ucsd.edu under MSV000086782 and ProteomeXchange under PXD023897 (http://proteomecentral.proteomexchange.org). The Spectronaut projects that are uploaded to the repositories can be viewed using the free Spectronaut viewer (www.biognosys.com/technology/spectronaut-viewer).

#### b. Exosome lipid cargo

All mass spectrometry raw files and datasets have been deposited to MassIVE ftp://massive.ucsd.edu/v07/MSV000094326/ under MSV000094326.

### Statistical analysis

Five biological replicates were analyzed for exosome plasma studies and four biological replicates for the senescent fibroblasts IMR90 studies. We used the two-tailed Student’s t-test or ANOVA (GraphPad Prism, La Jolla, CA, USA) to determine differences between the groups. Unless stated otherwise, the experimental data are presented as the mean ± standard deviation (SD) from three independent experiments.

### Pathway and network analysis

We conducted gene set over-representation analyses using the ConsensusPathDB-human tool, release 34 (15.01.2019). Specifically, we analyzed a list of all proteins that were significantly increased or decreased (q-value < 0.05) in (a) Old vs. young plasma, (b) IR vs. Qui, (c) doxo vs. Qui, and (d) MiDAS vs. Qui. We referenced curated pathways for enrichment analysis from the Gene Ontology (GO) Database, focusing on “Biological Process” term categories. We considered pathways with a minimum of three observed proteins, q-value < 0.01, and GO terms restricted to levels 4–5 as significant. To provide background reference, we used a list of all proteins present in the deep fractionated DDA spectral library. The complete list of differentially regulated pathways, corresponding proteins, reference pathway annotations, statistics, and source databases are available as a supplemental file (Table S4j, k). To generate dot plot representations of pathway analysis, we used the “ggplot2” package in R.

## Author Contributions

Conceptualization, BS, JC, ESB; experiments and methodology, SKP, JB, JRC, JPR, RLB, RB, LR; data processing, SKP, JB, JRC, RLB, LR; writing original draft, SKP, PYD, JRC; Figures, SKP, JRC, JB, BS; review and editing of the manuscript, BS, JB, JRC, ESB, MAW, CW, JBB, JC, LR.

## Acknowledgments

This work is supported by grants from the National Institutes of Health under award numbers U01 AG060906 (Schilling), U01 AG060906-02S1 and NCRR shared instrumentation grant S10 OD028654 (Schilling), U54 AG075932 (Schilling, Melov), P01 AG066591 (Ellerby), P01 AG017242 (Campisi) and R01 AG051729 (Campisi), P42 ES027704 (Baker), R01 GM141277 (Baker) and RM1 GM145416 (Baker). Dr. Patel was supported by a Glenn Fellowship in Aging Research from the Glenn Foundation. We thank John Carroll for providing graphical support and LC Sciences for miRNA Sequencing Services. Dr. Schilling and team were deeply saddened about the passing of Dr. Campisi and would like to recognize her as amazing scientist, mentor and friend.

