## Supplementary material for "Exosomes Released from Senescent Cells and Circulatory Exosomes Isolated from Human Plasma Reveal Aging-associated Proteomic and Lipid Signatures": Fig S1

### Supplementary Figures S1

**Fig. S1a** Workflow for the different plasma exosome enrichment methods were assessed for plasma exosome extraction.  SEC: size exclusion chromatography, UC: ultracentrifugation, UF: ultrafiltration**.**

**Fig. S1b** Bar graph showing the distribution of plasma exosome proteins identified using directDIA analysis in different exosome extraction methods UC and UF. UC: ultracentrifugation, UF: ultrafiltration

**Figure S1c:** Western blot confirms the presence of CD9, CD63, and TSG101 proteins (EVs markers) in cell culture extracted exosome, while the presence of Alpha-2-Macroglobulin (A2M) determined culture media protein contaminants.

**Figure S1a**

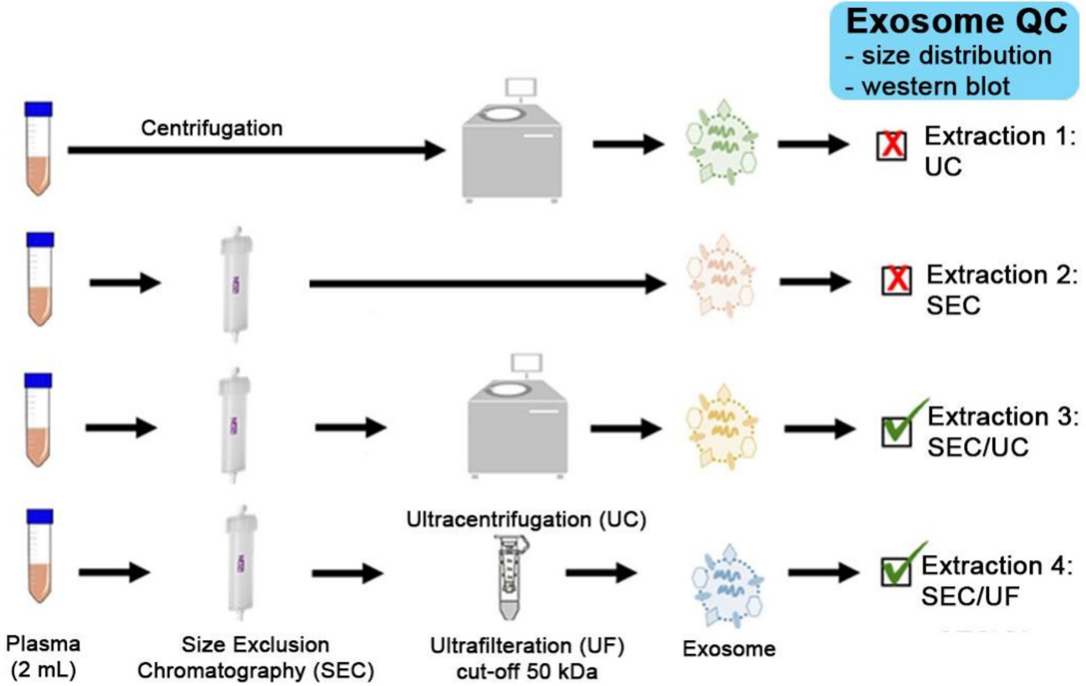

Exosome QC
- size distribution
- western blot
Centrifugation
Extraction 1:
UC
Extraction 2:
SEC
Extraction 3:
SEC/UC
Ultracentrifugation (UC)
Extraction 4:
SEC/UF
Plasma
(2 mL)
Size Exclusion
Chromatography (SEC)
Ultrafilteration (UF)
cut-off 50 kDa
Exosome

**Figure S1b**

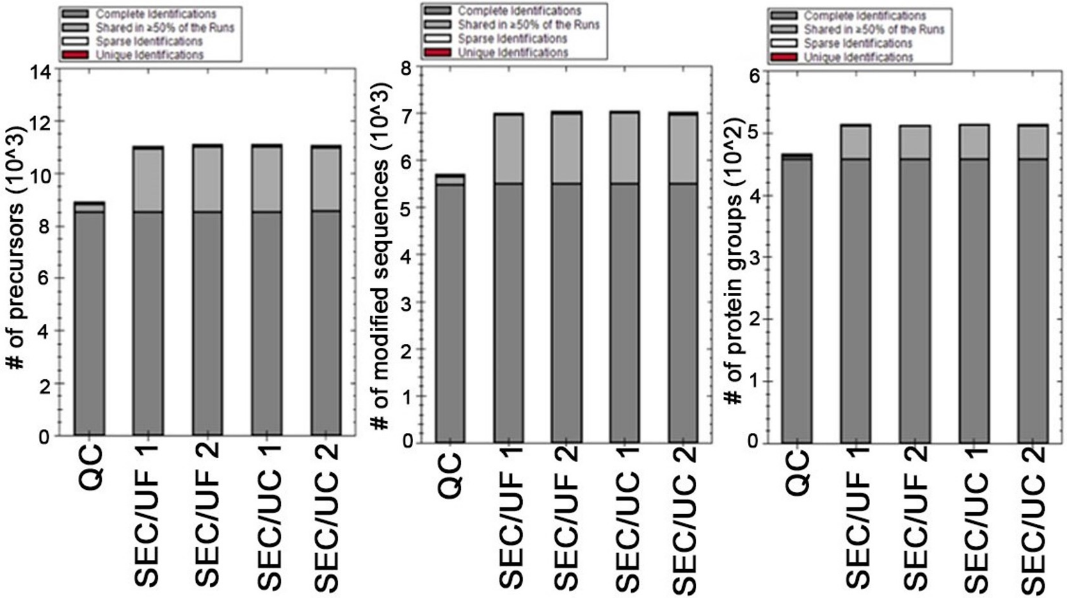


Protein identification
(≥2 peptides, Q value <0.01)

Total proteins = 578

**Figure S1c**

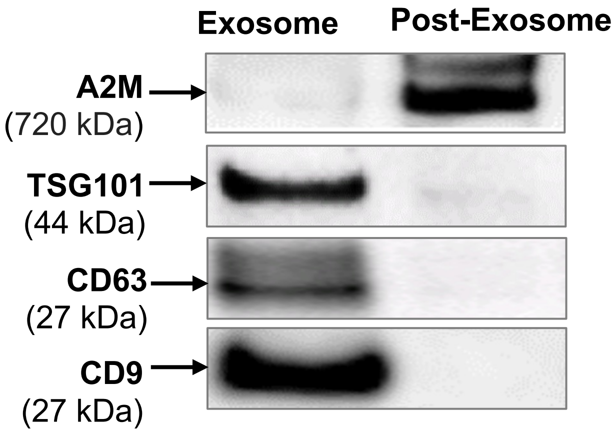

Exosome
Post-Exosome
A2M
(720 kDa)
TSG101
(44 kDa)
CD63
(27 kDa)
CD9
(27 kDa)
