## Supplementary material for "Exosomes Released from Senescent Cells and Circulatory Exosomes Isolated from Human Plasma Reveal Aging-associated Proteomic and Lipid Signatures": Fig S2

**Supplementary Figure S2**

**Figure S2a-d:** HPRP-fractionation and analysis of plasma exosomes for the generation of DDA, directDIA and hybrid-MS deep spectral proteomics.

**Fig. S2e Assessment of the DDA library.**
(i) Distribution of precursor m/z value. (ii) Distribution of precursor charge. (iii) Distribution of peptide length. (iv) Modifications of peptides. Carb (C), carbamidomethylation of cysteine residue; Oxi (M), oxidation of methionine residue; Ac (Prot N-ter), acetylation of protein N-terminus. (v) Number of peptides per protein group. (vi) Number of fragments per precursor. (vii) Fragment ion type. (viii) Fragment ion charge. (vi, vii, viii) Only fragments used in the assay were considered.

**Fig. S2f Assessment of the directDIA library.**
(i) Distribution of precursor m/z value. (ii) Distribution of precursor charge. (iii) Distribution of peptide length. (iv) Modifications of peptides. Carb (C), carbamidomethylation of cysteine residue; Oxi (M), oxidation of methionine residue; Ac (Prot N-ter), acetylation of protein N-terminus. (v) Number of peptides per protein group. (vi) Number of fragments per precursor. (vii) Fragment ion type. (viii) Fragment ion charge. (vi, vii, viii) Only fragments used in the assay were considered.

**Fig. S2g Assessment of the hybrid library.**
(i) Distribution of precursor m/z value. (ii) Distribution of precursor charge. (iii) Distribution of peptide length. (iv) Modifications of peptides. Carb (C), carbamidomethylation of cysteine residue; Oxi (M), oxidation of methionine residue; Ac (Prot N-ter), acetylation of protein N-terminus. (v) Number of peptides per protein group. (vi) Number of fragments per precursor. (vii) Fragment ion type. (viii) Fragment ion charge. (vi, vii, viii) Only fragments used in the assay were considered.

**Fig. S2h** Venn diagrams showing the common and unique protein groups in the plasma exosome libraries (DDA, directDIA and hybrid; ≥ 2 unique peptides) and the human ExoCarta database. Gene names were used to generate the Venn diagrams.

**Fig. S2i** Venn diagrams showing the common and unique exosome protein groups in the plasma exosomes libraries (DDA, directDIA and hybrid; ≥ 2 unique peptides) and the urine exosome library. UniProt IDs were used to generate the Venn diagrams.

### Figure S2

**a. Spectral library details**

| Sample Details | Database | Precursors | Peptides | Proteins | Protein groups |
|---|---|---|---|---|---|
| 25 DDA | HPRP fractionation dataset (DDA) | 43,201 | 26,655 | 5,186 | 2,323 |
| 10 directDIA | directDIA dataset (DIA) | 21,944 | 12,357 | 1,787 | 856 |
| 25 DDA + 10 directDIA | Hybrid dataset (DDA + DIA) | 44,981 | 26,172 | 4,993 | 2,225 |

**b. Peptides**

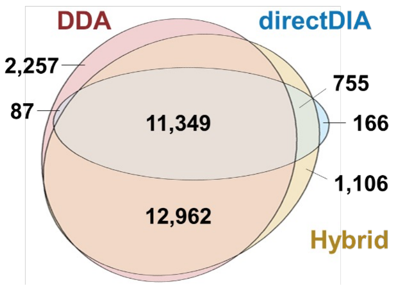


**c. Protein groups**

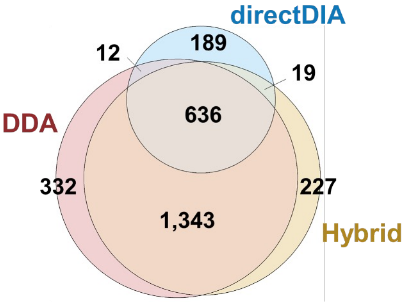


**d. QC analysis of library**

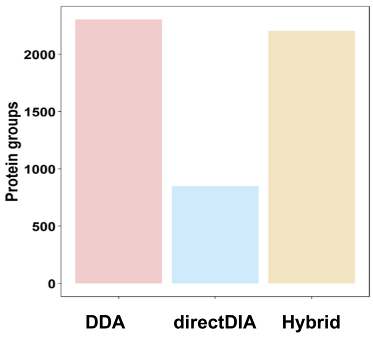

**Figure S2e: Assessment of the DDA library.**

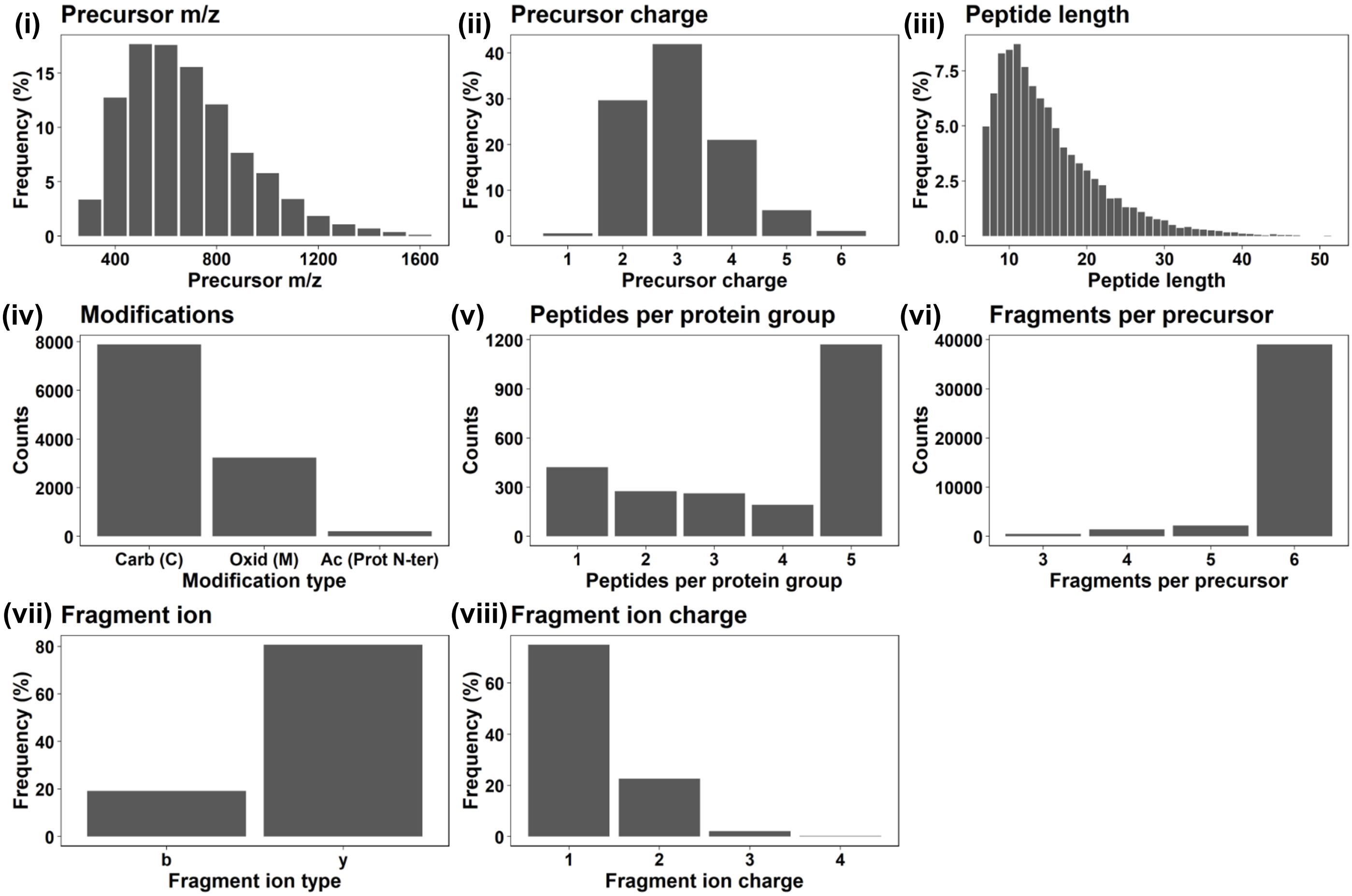

**Figure S2f: Assessment of the directDIA library.**

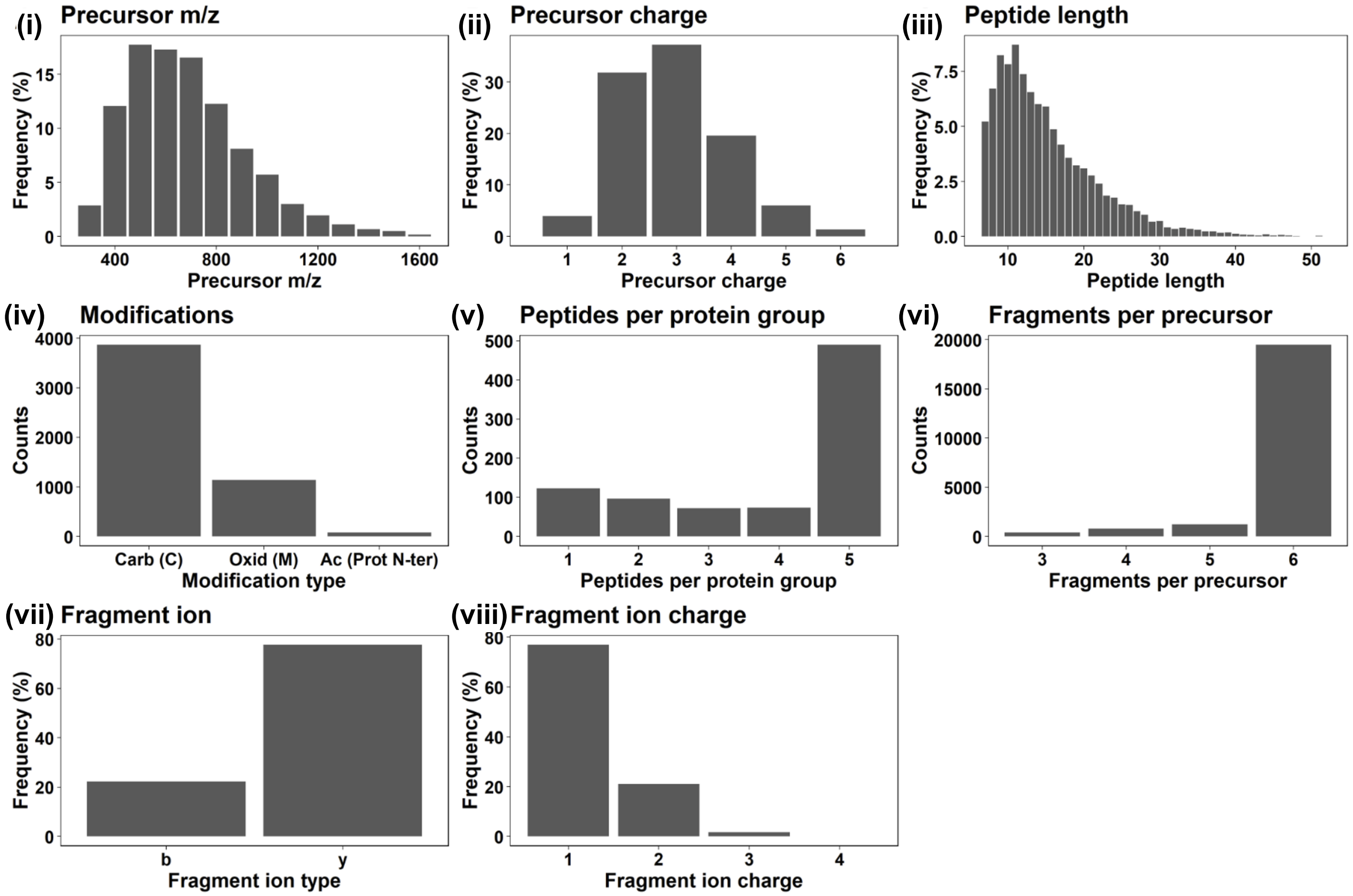

Figure S2g: Assessment of the hybrid library.

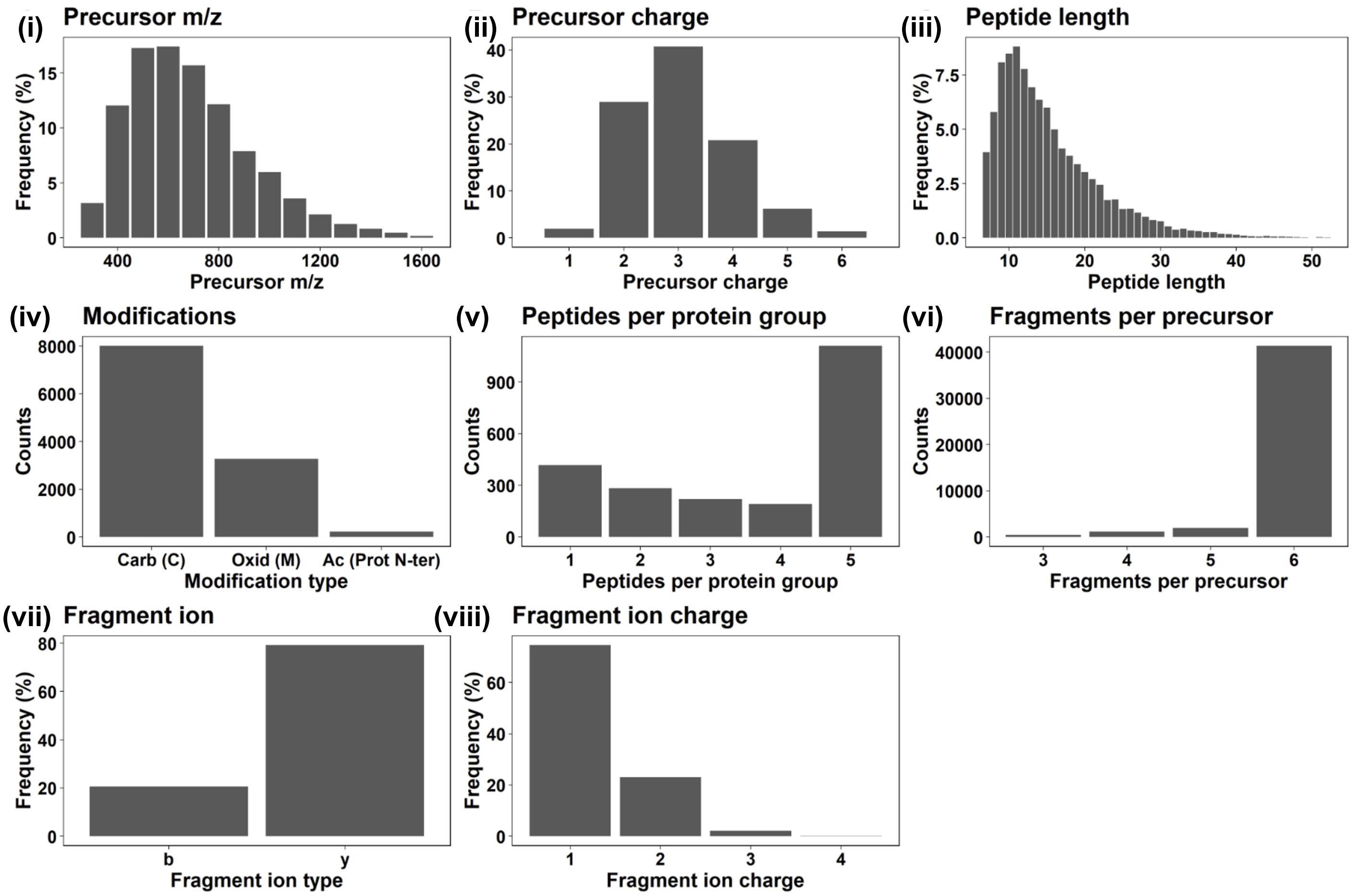

**Figure S2h**

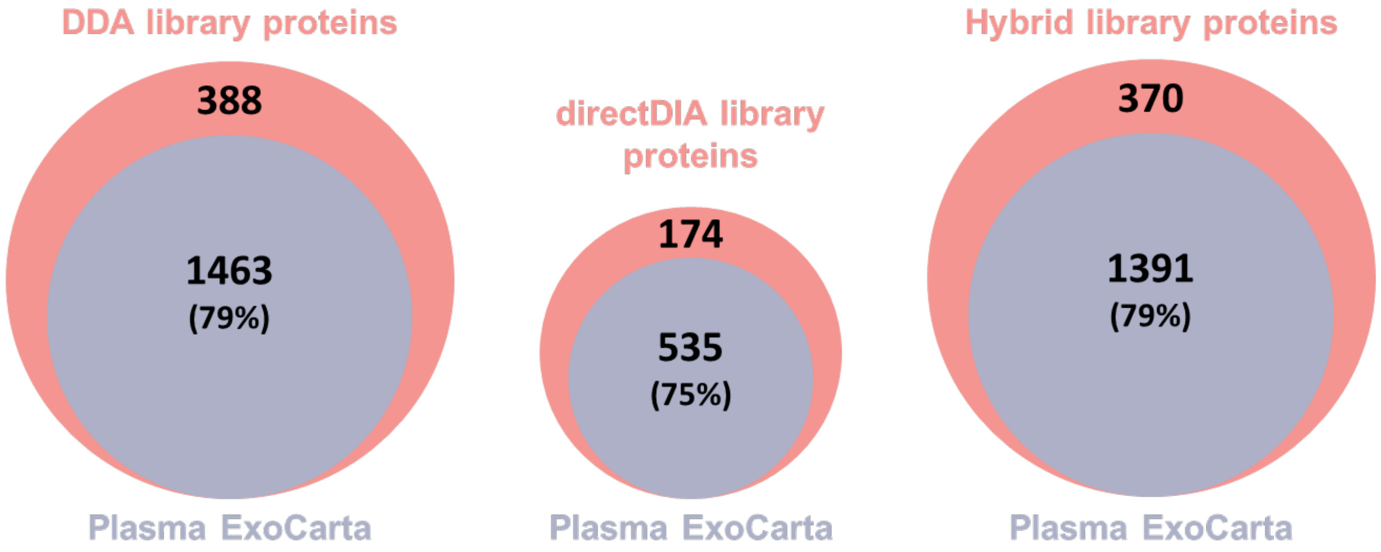


**Figure S2i**

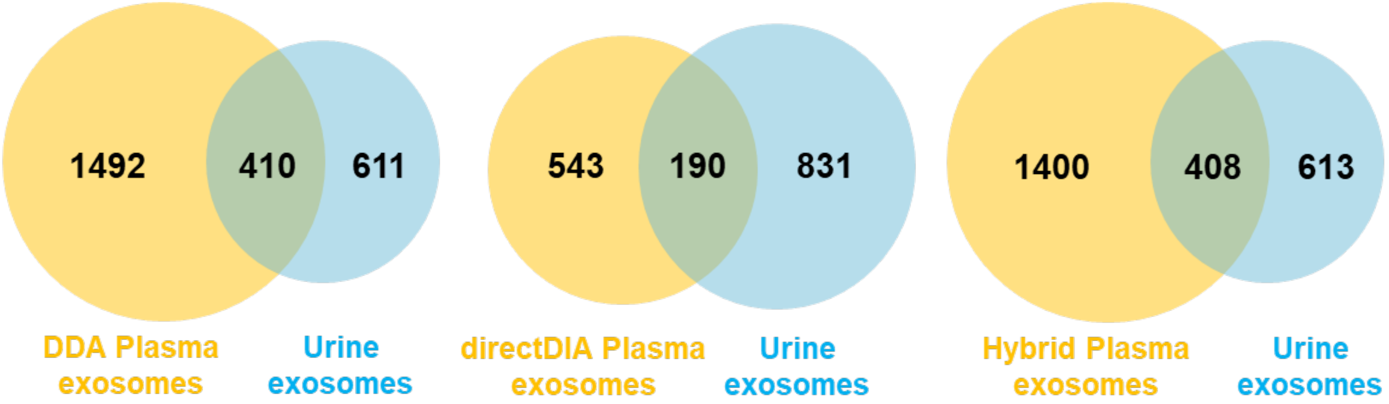
