## Supplementary material for "Exosomes Released from Senescent Cells and Circulatory Exosomes Isolated from Human Plasma Reveal Aging-associated Proteomic and Lipid Signatures": Fig S3

### Supplementary Figure S3

**Figure S3a:** Estimation of cell death using SYTOX Green Cytotoxicity Assay

**Figure S3b:** NAD+/NADH ratio calculations for IMR90 cells

**Figure S3c:** Volcano plots showing the significantly altered protein groups in senescence (IR, Doxo, MiDAS) vs Quiescent (control) out of the 1,426 quantifiable protein groups (q-value ≤ 0.05, $|log_2$fold change$| \geq 0.58$)

Figure S3a

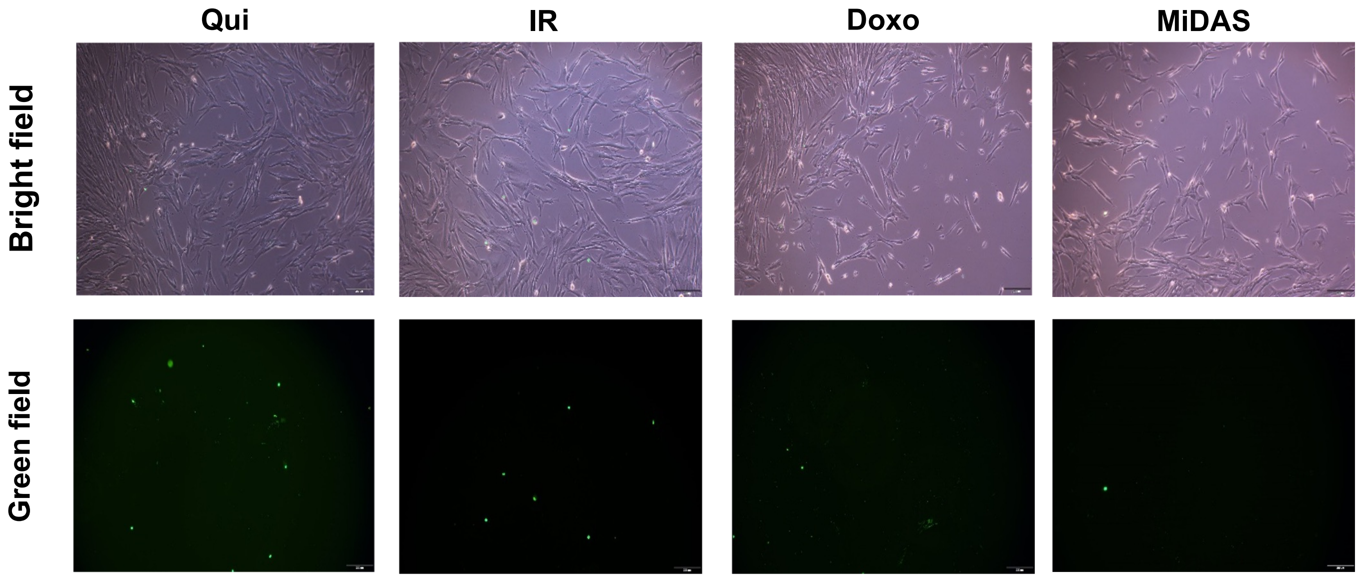

Figure S3b

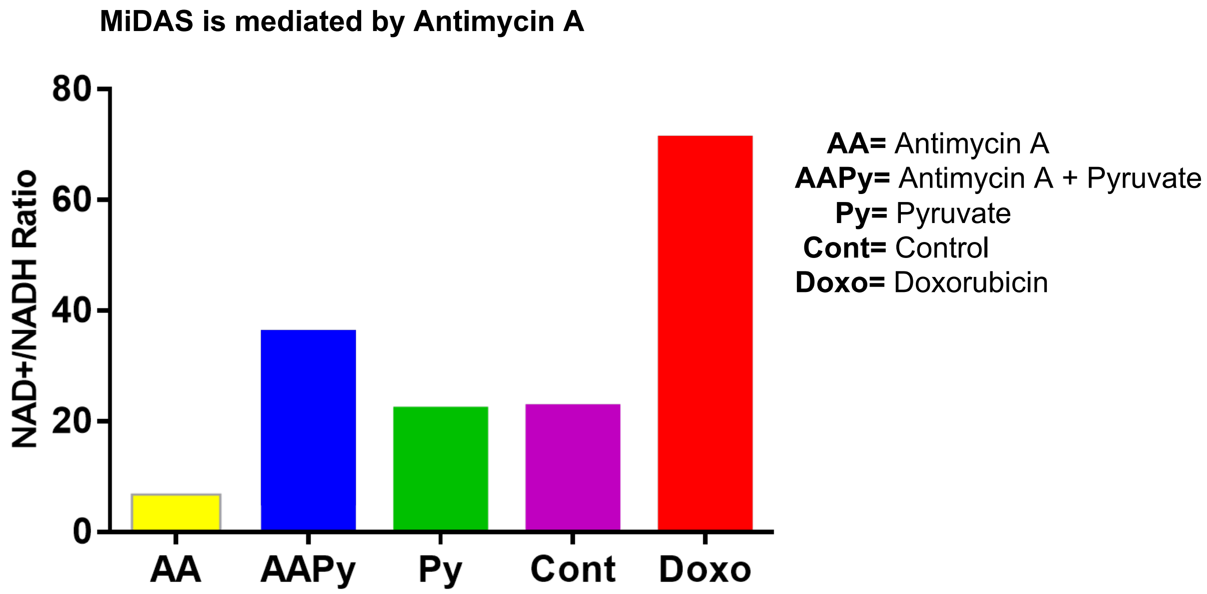
MiDAS is mediated by Antimycin A
NAD+/NADH Ratio
80
60
40
20
0
AA
AAPy
Py
Cont
Doxo
AA= Antimycin A
AAPy= Antimycin A + Pyruvate
Py= Pyruvate
Cont= Control
Doxo= Doxorubicin

### Figure S3c

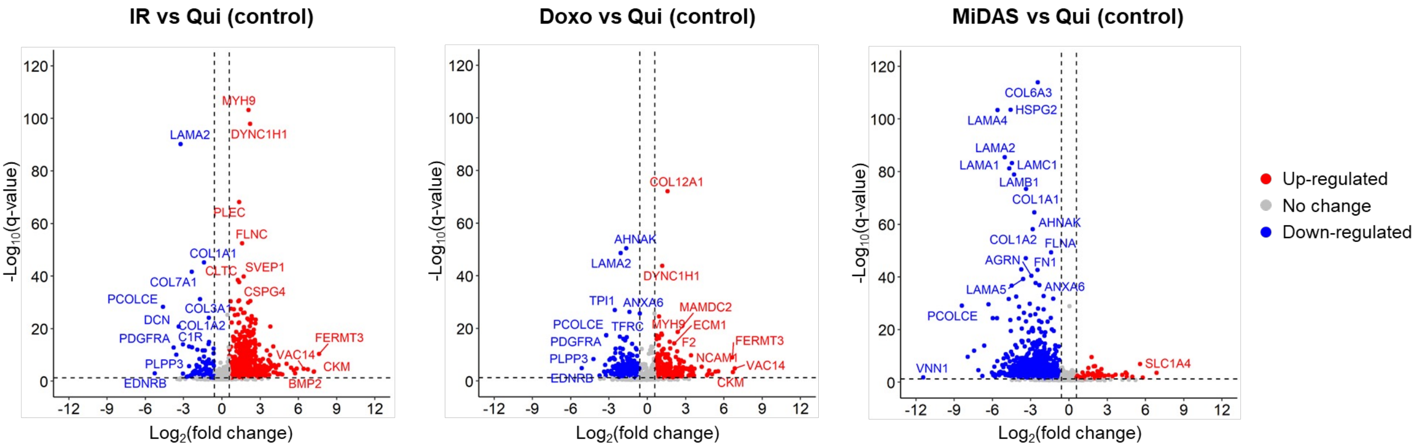

IR vs Qui (control)
Doxo vs Qui (control)
MiDAS vs Qui (control)
-Log10(q-value)
Log2(fold change)
MYH9
DYNC1H1
LAMA2
PLEC
FLNC
COL1A1
CLTC
SVEP1
COL7A1
CSPG4
PCOLCE
COL3A1
DCN
COL1A2
C1R
PDGFRA
FERMT3
VAC14
CKM
PLPP3
EDNRB
BMP2
COL12A1
AHNAK
DYNC1H1
TPI1
ANXA6
MAMDC2
TFRC
MYH9
ECM1
F2
NCAM1
COL6A3
HSPG2
LAMA4
LAMC1
LAMB1
LAMA1
FLNA
AGRN
FN1
LAMA5
VNN1
SLC1A4
Up-regulated
No change
Down-regulated
