## Supplementary material for "Exosomes Released from Senescent Cells and Circulatory Exosomes Isolated from Human Plasma Reveal Aging-associated Proteomic and Lipid Signatures": Fig S4

### Supplementary Figure S4

**Figure S4a-c:** Bar diagrams showing the total number of identified and quantified protein groups in the human plasma exosome study with the three different libraries (hybrid, DDA, and directDIA).

**Figure S4d:** Venn diagram showing the common and unique significantly altered human plasma exosome protein groups obtained with the three different libraries

**Figure S4e:** Boxplot showing the ratio of the ‘Old vs Young’ fold-change for the shared significantly altered protein groups obtained with directDIA or hybrid library vs DDA library

**Figure S4f:** Summary of all quantifiable (with ≥ 2 unique peptides) and significantly altered age-specific plasma exosome protein groups (q-value < 0.05, $|\log_2$ fold change$| \geq 0.58$)

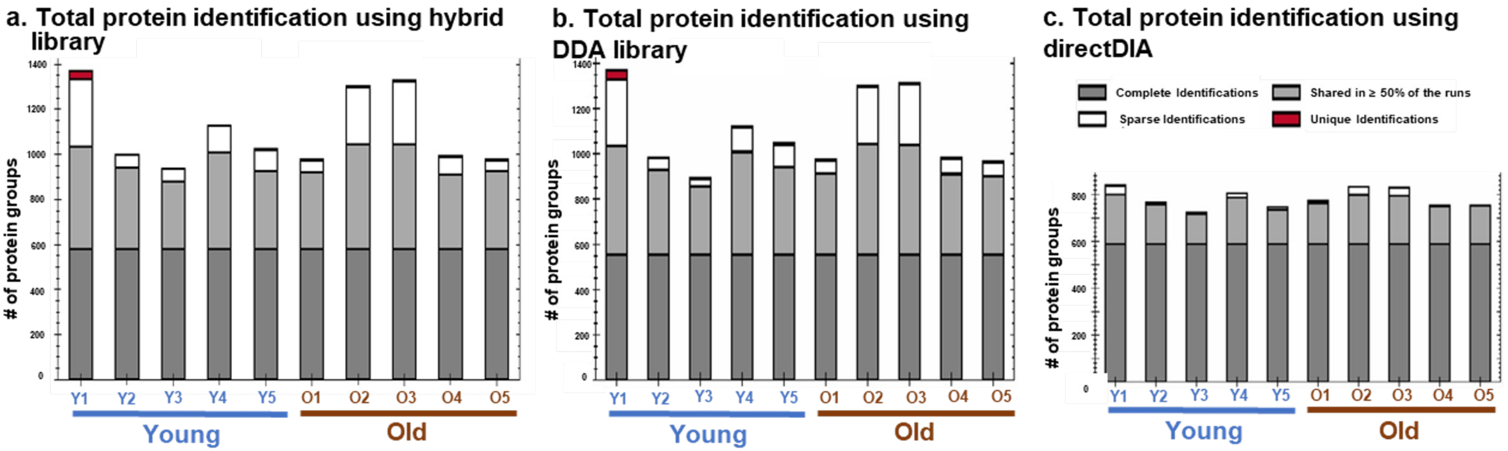


We used DIA-MS protein identification and quantification to analyze plasma exosomes from young and elderly individuals and processed the data using three different spectral libraries: hybrid DDA-DIA library, DDA library, or directDIA library. Overall, we reproducibly identified and quantified a total of 1,356 protein groups using the hybrid spectral library (Fig. S4a, Table S4B), 1,349 protein groups using the spectral library generated from DDA searches (Fig. S4b, Table S4E), and 760 protein groups when applying directDIA searches (Fig. S4c, Table S4H). For all, we reported protein groups numbers obtained with a 1% false discovery rate (FDR) and ≥ 2 unique peptides.

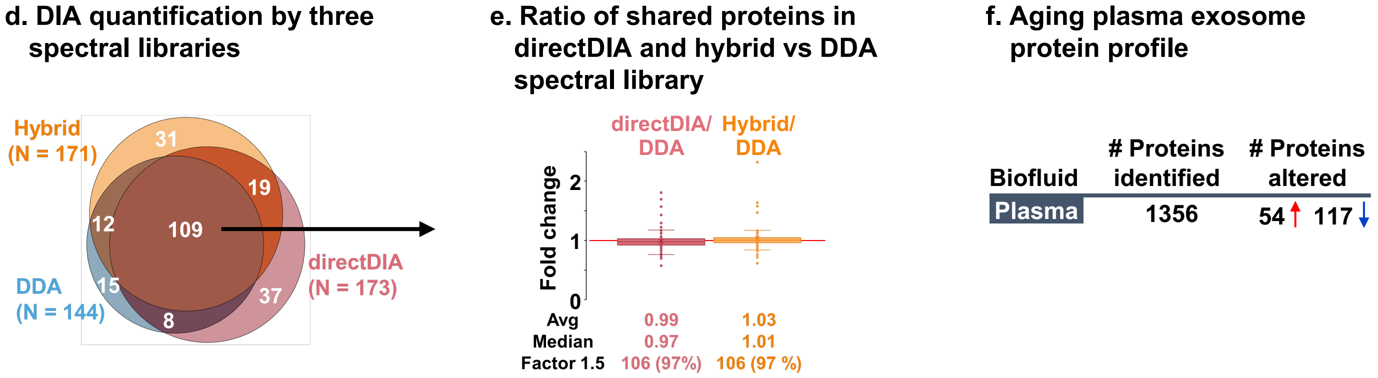


We investigated the significantly changing protein groups in plasma exosomes from ‘old vs young’ individuals that resulted from all spectral library-based approaches, and interestingly 109 significantly altered proteins were commonly shared between all libraries (q-value < 0.05 and $|\log_2$ old vs young$| \geq 0.58$; Fig. S4d). The highly consistent fold-changes of the shared protein groups with the hybrid and directDIA library compared to the DDA spectral library demonstrate that all spectral library workflows are robust and can be used for protein quantification (Fig. S4e). However, the maximum depth of coverage was obtained with the hybrid spectral library with 1,356 identified and quantified protein groups (≥ 2 unique peptides), so we focused on the 171 protein groups that significantly changed within plasma exosomes between the ‘old vs young’ cohorts, when using the DIA-MS hybrid library approach (Fig. S4f), in this study.
