## Supplementary material for "Exosomes Released from Senescent Cells and Circulatory Exosomes Isolated from Human Plasma Reveal Aging-associated Proteomic and Lipid Signatures": Fig S5

**Figure S5. Age-specific plasma exosome miRNA signatures in old and young cohorts.**

**A.** miRNA nucleotide size distribution. **B.** Overlapping and unique exosome miRNAs in plasma from older and young individuals. **C.** Volcano plot showing significantly altered age-specific plasma exosome miRNAs (P-value < 0.05 and > 1.5-fold). Red, upregulated; Green, down-regulated; and Gray, not significantly changed. A few differentially abundant miRNAs are labeled. **D.** Age-related exosome miRNA pathways.

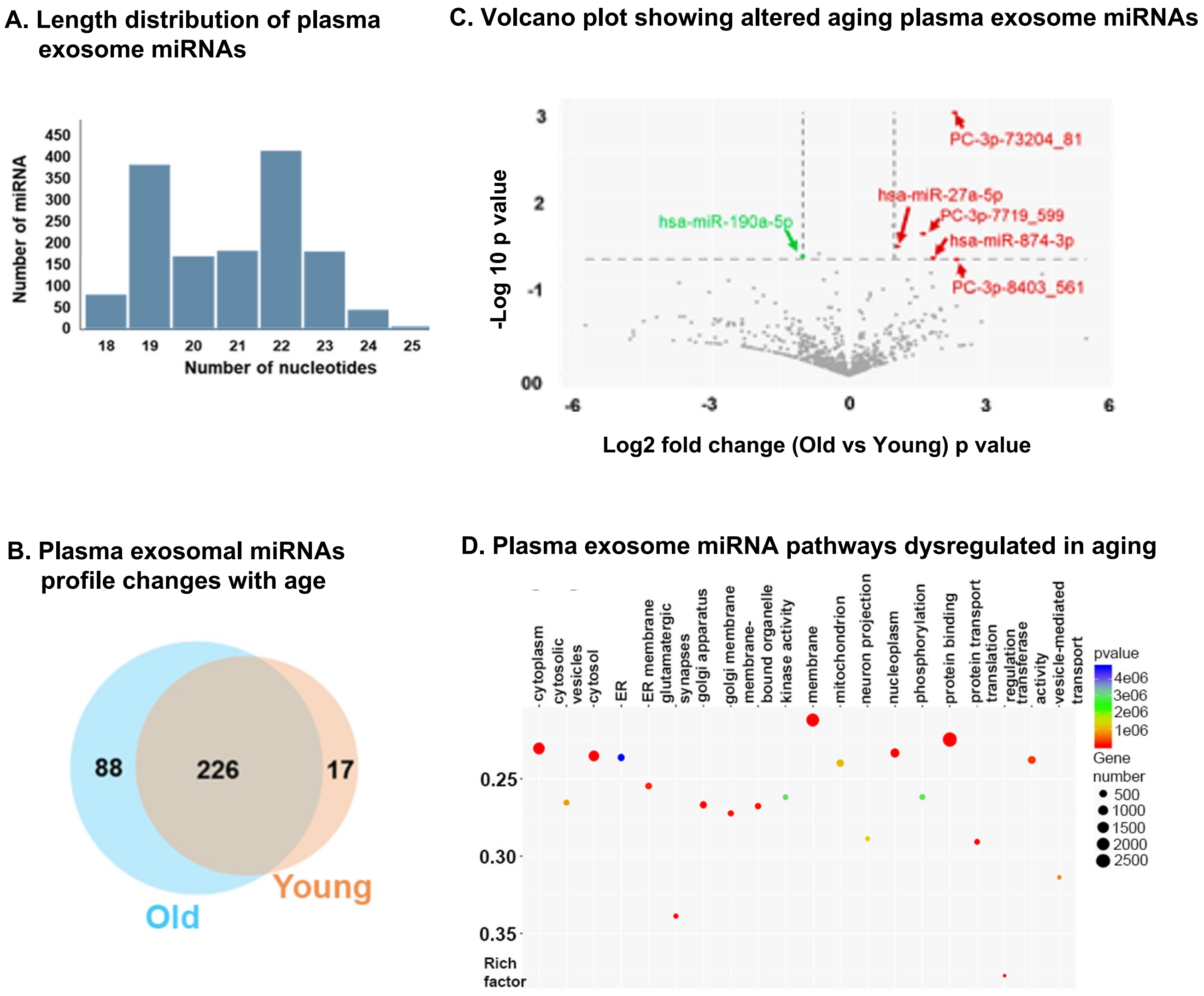
