## Supplementary material for "Exosomes Released from Senescent Cells and Circulatory Exosomes Isolated from Human Plasma Reveal Aging-associated Proteomic and Lipid Signatures": Fig S6

### Supplementary Figure S6

Venn diagram showing the common and unique significantly altered protein groups in human plasma exosomes from this study (old vs young comparison, “Plasma exosome proteins”), in senescence-induced IMR90 exosomes (merged from the three inducers) from this study (“Exosome SASP proteins”), and human age-associated soluble plasma proteins from Johnson, Wyss-Coray *et al.*, 2020 (“Soluble plasma proteins”) [32].

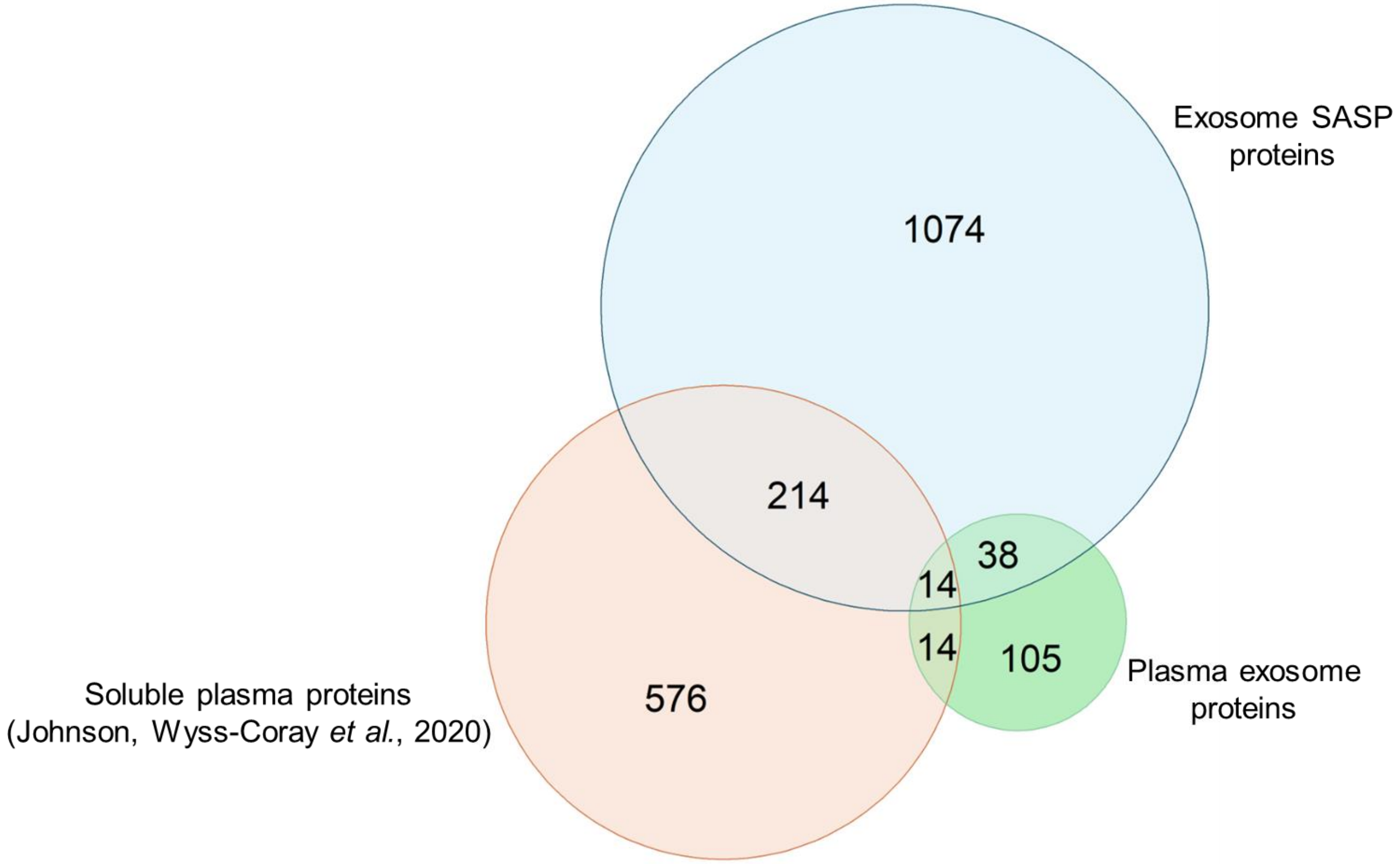
